# Local minimization of prediction errors drives learning of invariant object representations in a generative network model of visual perception

**DOI:** 10.1101/2022.07.18.500392

**Authors:** Matthias Brucklacher, Sander M. Bohte, Jorge F. Mejias, Cyriel M. A. Pennartz

## Abstract

The ventral visual processing hierarchy of the cortex needs to fulfill at least two key functions: Perceived objects must be mapped to high-level representations invariantly of the precise viewing conditions, and a generative model must be learned that allows, for instance, to fill in occluded information guided by visual experience. Here, we show how a multilayered predictive coding network can learn to recognize objects from the bottom up and to generate specific representations via a top-down pathway through a single learning rule: the local minimization of prediction errors. Trained on sequences of continuously transformed objects, neurons in the highest network area become tuned to object identity invariant of precise position, comparable to inferotemporal neurons in macaques. Drawing on this, the dynamic properties of invariant object representations reproduce experimentally observed hierarchies of timescales from low to high levels of the ventral processing stream. The predicted faster decorrelation of error-neuron activity compared to representation neurons is of relevance for the experimental search for neural correlates of prediction errors. Lastly, the generative capacity of the network is confirmed by reconstructing specific object images, robust to partial occlusion of the inputs. By learning invariance from temporal continuity within a generative model, despite little change in architecture and learning rule compared to static input- reconstructing Hebbian predictive coding networks, simply by shifting the training paradigm to dynamic inputs, the approach generalizes the predictive coding framework to dynamic inputs in a more biologically plausible way than self-supervised networks with non-local error-backpropagation.

**Author Summary:** Neurons in the inferotemporal cortex of primates respond to images of complex objects independent of position, rotational angle, or size. While feedforward models of visual perception such as deep neural networks can explain this, they fail to account for the use of top-down information, for example when sensory evidence is scarce. Here, we address the question of how the neuronal networks in the brain learn both bottom-up and top-down processing without labels as they are used in the artificial supervised learning paradigm. Building on previous work that explains vision as a process of iteratively improving predictions, learning in the predictive coding network is driven by the local minimization of prediction errors. When trained on sequences of moving inputs, the network learns both invariant high-level representations comparable to those in the inferotemporal cortex of primates, and a generative model capable of reconstructing whole objects from partially occluded input images in agreement with experimental recordings from early visual areas. Advancing the search for experimental hallmarks of prediction errors, we find that error neurons in the higher areas of the network change their activity on a shorter timescale than representation neurons.

## 1. Introduction

How networks of neurons in the brain infer the identity of objects from limited sensory information is one of the preeminent questions of neurobiology. Strengthening theories of generative perception (1–5), evidence has accumulated to suggest that the mammalian perceptual system is relying on various forms of prediction to facilitate this process. Across time, repetition suppression that requires explicit expectations (6, 7), encoding of deviation from temporal expectations in macaque’s inferotemporal and prefrontal cortex (8, 9) and encoding of expected movement outcomes in mouse V1 (10) show that the brain constantly tries to predict future inputs. V1 activity evoked by illusory contours (11, 12), encoding of information from occluded scene areas in early visual areas of humans (13) and modulation of neural responses by expectations based on the surrounding context (14) show that predictions are not only made forward in time, but also across space (in the present). According to predictive coding theory, these predictions are mediated by corticocortical top- down connections (3) and then corrected based on the received bottom-up input (5) in line with hierarchical Bayesian perception (15). Predictive coding models have successfully explained properties of the visual system such as end-stopping in V1 neurons and learning of wavelet-like receptive fields (5) and V1 activity in illusory contours (16, 17). However, these studies are focused on low-level effects, while the learned higher-level representations have been investigated much less (although see (18) for learning of sparse representations).

Continuously generated by the awake brain, neural representations of the external world form a partial solution to the problem of inference, arguably constituting the basis of conscious experience (19), decision-making and adaptive planning (20). They can be loosely defined as activity patterns in response to a sensory stimulation elicited by an object. Especially important is the ability to represent multiple views of the same object in similar patterns of activity. These invariant representations have two key advantages: First, information acquired about an object (such as a novel action associated with it) can be linked to only one representation, making learning more efficient. Secondly, as illustrated in Fig 1, the newly acquired invariant information about single objects generalizes automatically across all viewing conditions, facilitating learning from few examples. Evidence for invariant neural representations comes from the ventral temporal lobe (21), the hippocampus in humans (22), inferotemporal cortex of rhesus (23, 24) and macaque monkeys (25) as well as rats’ laterolateral extrastriate area (LL) (26, 27). Current theories of how neurons come to acquire such a specialized tuning either fail to account for fundamental aspects of brain circuitry and physiology or rely on artificial learning paradigms. To construct useful representations, biological systems are limited to mostly unsupervised learning (from unlabeled data) and local learning rules, whereas machine vision algorithms based on neural networks typically rely on large amounts of labeled training data and use mechanisms like weight-sharing (28). These mechanisms facilitate generalization across viewing conditions but lack a biological foundation.

**Fig 1.**
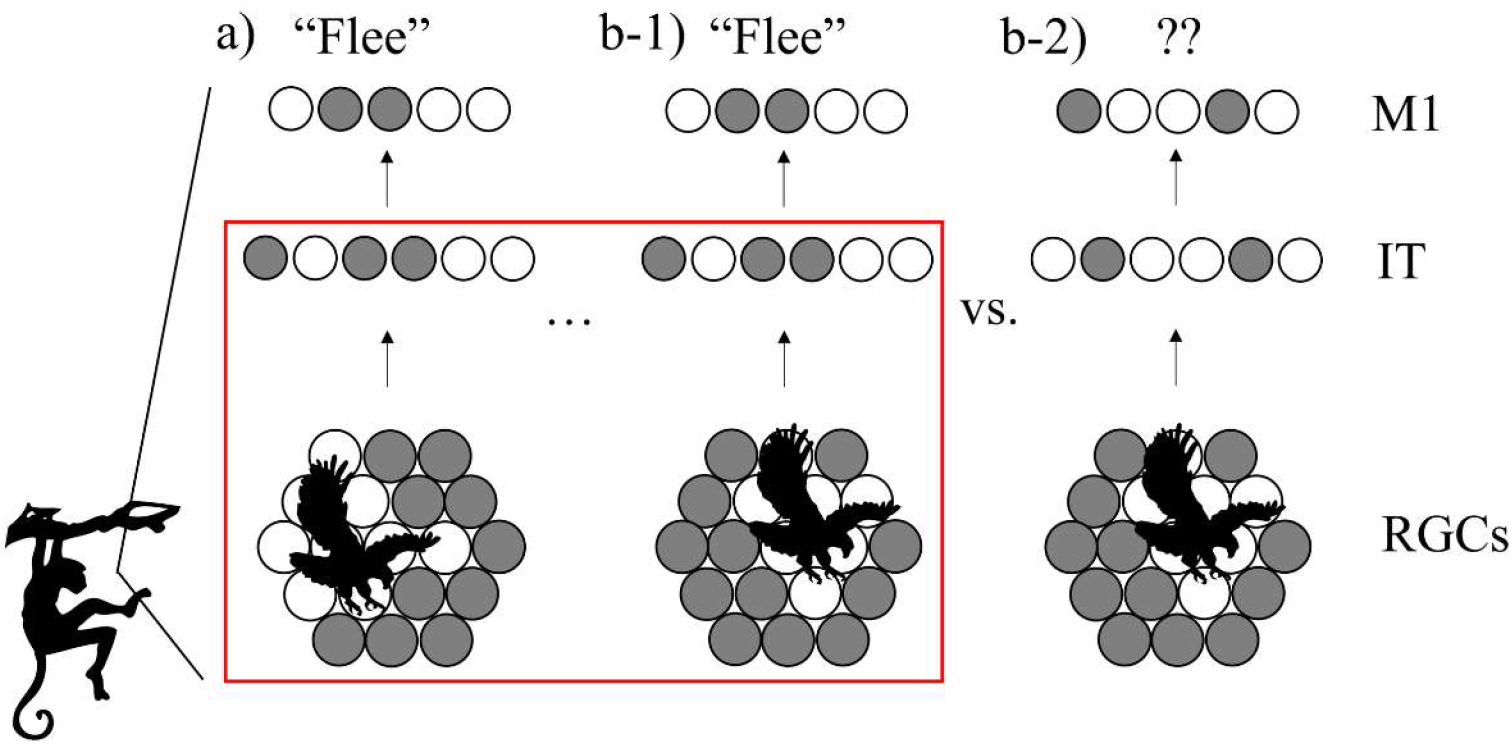
View-invariant representations for efficient cognition. a) Barely escaping an attack, the monkey learns to associate an action (”flee”, encoded by the neural pattern in primary motor area M1) with the activity pattern in its retinal ganglion cells (RGCs, bottom) triggered by the image of an approaching eagle. Active cells are shown in grey. b-1) When the monkey later encounters a similar eagle from a different angle, an invariant higher-level representation (center) can still trigger the same action. b-2) Without invariant coding, the action does not generalize to this viewing condition. Red box: Scope of this paper: How do multiple low-level activity patterns become linked to one high-level, invariant representation?

A biologically plausible approach to learn view-invariance from transformation sequences is so-called trace learning (29–31) which is linked to Slow Feature Analysis (SFA) (32). It is based on the idea that temporal proximity between sensory patterns should be reflected in representational similarity, as the assumption can be made about the world that the causes (objects etc.) vary more slowly than the stimulation patterns they evoke on the retina. Indeed there is evidence for the importance of temporal stimulus continuity for learning of transformation-tolerance in early visual areas of rats (33) and area IT of monkeys (34). Based on this principle of representing consecutive inputs similarly, Halvagal and Zenke (35) recently showed that a more intricate learning rule with additional variance maximization leads to disentangled high-level representations. Other self-supervised models avoid representational collapse through contrasting examples (36).

However, all of these models process information in a strictly feedforward manner or limit the role of feedback connections to a modulatory function, in contrast to evidence on retinotopic, content-carrying feedback connections in the visual cortex (37–39). Here, we propose a common underlying learning mechanism for both high-level representations and a generative model capable of reconstructing specific of sensory inputs: the minimization of local prediction errors through inference and learning.

Like the abovementioned feedforward models of invariance learning, predictive coding offers a mechanism for maintenance of higher-level representations: they are only updated when lower levels send up error signals. It can be implemented in a hierarchical neural network model of the visual processing stream using local, Hebbian learning. Furthermore, it is intimately related to the abovementioned slowness principle, which states that the most meaningful features often change on a slow timescale (40), because extracted causes tend to be good predictors for future input (41). To sum up, predictive coding is a promising candidate to explain learning of invariant object representations within the framework of generative modelling.

To acquire transformation-tolerance from temporal continuity, input sequences are required. Most predictive coding models so far, however, either operate on static inputs (5,18,42) or use non-local learning rules (43) such as backpropagation (44, 45) and biologically implausible LSTM units (16, 46). Here, we train multilayered predictive coding networks with little architectural modifications from (5, 18) on transformation sequences with purely Hebbian learning. We confirm learning of a generative model, showing that top-down predictions made by the network approximate the original input. Importantly, these predictions are not forward in time, but across retinotopic space, representing the current input. Presented with partially occluded input sequences, the network pattern-completes the occluded areas through top- down feedback, mimicking functions of human V1 and V2. While reconstructions from lower areas are more faithful, predictive neurons in the network’s higher areas develop view- invariant representations akin to responses of neurons in the inferotemporal area of primate cortex: Input stimuli shown in temporal proximity are represented similarly. A decoding analysis confirms that distinct objects are well separable. Lastly, the temporal dynamics of the neural subpopulations are analyzed and compared to recent electrophysiological data from rats. As in the experiment, temporal stability of representation neurons (measured by the decay of autocorrelation) increases as one moves up the hierarchy. In addition, the model makes the prediction that high-level error-coding neurons operate on a faster timescale than their representational counterparts.

## 2. Methods

We developed a neural network consisting of four hierarchically arranged areas. Applying the principles of predictive computation, we restricted ourselves to the minimally necessary components, but other connectivity patterns are conceivable (suggested e.g. by Heeger (47)). As in previous implementations of predictive coding (5, 18), each area contains two subpopulations of neurons that are illustrated in Fig 2 :

1. Representation neurons collectively hold the ”inferred causes”, in higher areas corresponding to perceptual content. Together with the synaptic connections towards lower areas, they generate top-down predictions to match the current representations in the area below.
2. Error neurons measure mismatch between representation neuron activity (in the lowest area: the sensory input) and top-down predictions.
3. Some models such as (48) suggest computation of errors in dendrites, but based on the evidence for neural encoding of errors (37), we assign dedicated neurons to encode them. Development of such error-tuned neurons has been modelled by (49) in cortical microcircuits and by (50) as a result of energy efficiency. The number of neurons in each area depends on the dimensions of the dataset and is given in section 6.1 of the Supplementary Material, supported by an analysis of how altering the number of neurons affects decoding performance in Supplementary Material 6.13.

**Fig 2.**
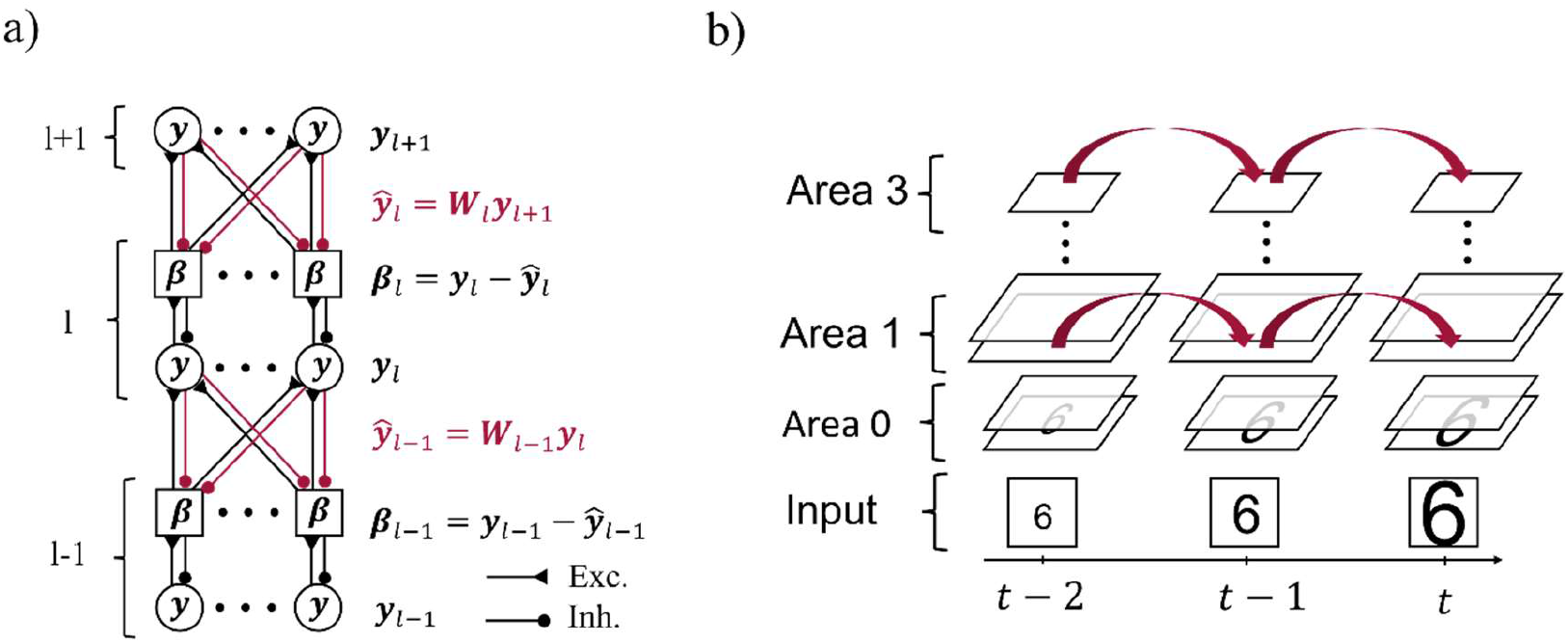
Model architecture and inference on video sequences. a) Representational activity ***y****_l_* in area l (l ∈ {1,2}) is influenced by both top-down predictions highlighted in red via the respective top-down errors *β_l_*, and by the bottom-up errors *β_l_*_-1_. Intra-area connections between representation neurons (circles) and respective error neurons (squares) are one-to-one, while inter-area connections are all-to-all. Synaptic connections between neurons are drawn as filled circles if inhibitory and as triangles if excitatory. b) A sequence of input images is fed into the lowest area of the network across subsequent moments in time(*t* − 2, …, *t*). The network maintains representational activity through time and thus uses it as a prior for the inference of subsequent representations.

### 2.1 Inference: Updating neural activity

At the start of a sequence, all neural activity is set to a uniform, low value (unless stated differently in the Results section). While an image is presented to the network, the lowest area representation neurons linearly reflect the pixel-wise intensity of the input (at the bottom of Fig 2b). Error neurons in area *l* receive excitatory input from the activity ***y****_l_* of associated representation neurons as shown in the one-to-one connections in Fig 2a, and are inhibited by the summed-up predictions 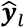 from the higher area:

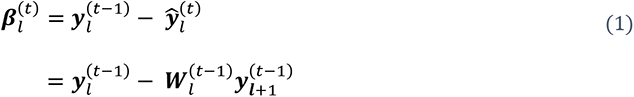

where bold letters indicate vectors and matrices and 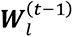 denotes the symmetric weight matrix between area l and area l+1 from the previous time step (the weights will change during learning). Strictly symmetric weight matrices as frequently used in predictive coding models (5, 18) lead to a weight transport problem during learning. However, it has been shown that, in combination with weight decay, symmetric weights can be obtained by learning rule comparable to ours without explicitly enforcing symmetry (51), since the locally available pre- and postsynaptic activity that determine the weight change are identical (symmetric) for each pair of feedforward and feedback connections. Each representation neuron receives inhibitory input from one error neuron in the same area and excitatory input from the weighted bottom-up errors and thus changes its activation state at each time step (see “inference” in Alg S1:

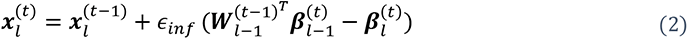

This adjustment of neural activation state ***x****_l_* (akin to membrane potential) of representation neurons can be interpreted as matching top-down predictions better than before (and thus reducing activity of the associated error neuron) and sending down predictions that better match representation neuron activity in the area below (thus reducing errors there). The rate at which neuronal activation is changed is governed by the parameter *ɛ_inf_* referred to in the following as the inference rate. The activation state ***x****_l_* is now translated into an output firing rate ***y***_1_ :

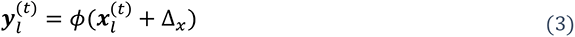

where ϕ denotes the sigmoid activation function, and Δ*_x_* a constant lateral offset of the firing threshold. The saturation of the sigmoid for large inputs corresponds to a maximal firing rate of the representation neurons, in contrast to the more artificial, (rectified) linear activation functions used in (5) and (18) that do not have an upper bound.

### 2.2 Learning without labels: Updating synaptic strengths

Before training, weights are initialized to random values from a Gaussian distribution centered at zero and with standard deviation of 0.5, clipped at zero to prevent negative weights and divided by the number of neurons in the next (higher) area. After a fixed number of inference steps (Supplementary Material 6.1), long-term adaptation of synaptic weights is conducted in a Hebbian manner, strengthening synapses between active error neurons in area l and simultaneously active representation neurons in the area above (l+1):

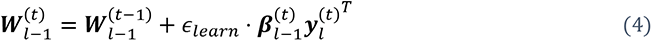

with learning rate *ɛ_learn_*. Apart from not using weight decay or normalization, we thus use the same learning rule as (5, 18). Based on the slower change of synaptic efficacy in comparison to membrane potential dynamics, weights are assumed to be constant between these updates. In Eq 4, the sign of the prediction error controls the direction of the weight change. If the prediction is too large relative to the activity of the representation neurons in this area, the error is negative, and the weight mediating the prediction will be reduced. As a result, given the same prediction, the error in the consecutive time step will be smaller. This stabilizing effect on the response of error neurons is familiar from the work of (52) that showed how Hebbian plasticity regulates inhibitory input to reduce firing and achieve a balanced global state.

To summarize, both the balanced, excitatory-inhibitory wiring of the network and the unsupervised adaptation of weights based on remaining prediction errors lead to an alignment of representations and predictions, and thus a reduction in error neuron activity. The sum of squared prediction errors can then be seen as an implicit objective function, upon which the inference steps conduct an approximate gradient descent (taking into account only the sign and not value of the derivative of the activation function unlike (53)), and upon which learning conducts a precise gradient descent.

### 2.3 Training procedure

We trained the network on temporally dynamic inputs, using short video sequences. After validating network performance on moving horizontal and vertical bars, we switched to using ten digits of the MNIST handwritten digits dataset (one per digit from 0-9). Each sequence contained six gradually transformed images, and separate datasets were created for translational motion, rotation, and scaling (Fig 3). For translational and rotational motion, two transformation speeds were used, differing in overlap between consecutive images. The examples shown in Fig 3 are from the dataset with larger step size (“fast” condition). To further examine robustness of the training paradigm under more realistic and less sparse inputs, random noise patterns were added to the image background during training. A last dataset consisted of five high-pass filtered images of toy objects (an airplane shown in the last row of Fig 3, a sports car, a truck, a lion and a tin man) from the smallNORB dataset (54), undergoing a rotation.

**Fig 3.**
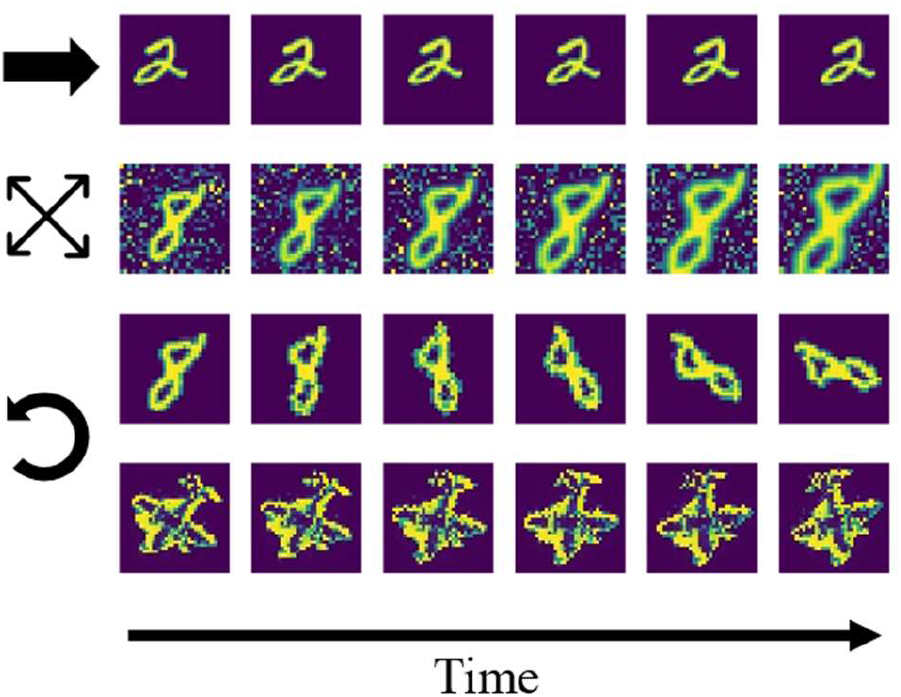
Example sequences from the stimulus datasets. First row: digit translation without noise, second row: digit scaling with noise, third and fourth row: rotation of digit/toy plane without noise.

The network was trained on the ten (for the toy objects: five) sequences, each presenting a different digit, for multiple epochs. Each epoch consisted of iterations of the same sequence (e.g. of a moving digit ’6’) before switching to the next (of digit ’7’). Hyperparameters such as the number of repetitions are listed in Supplementary Material 6.1. This repetition of individual sequences drastically improved network performance and could be achieved by the brain through a replay or reactivation mechanism (observed in visual cortex by (55, 56), see also (57, 58)). For laterally moving stimuli, repeated presentation can also be achieved by object-tracking saccades that lead to repeated motion across the same photoreceptors on the retina. As the most information-neutral state, the activity was reset to uniform, low values at the beginning of each sequence. This assumption is justified for objects that are seen independently of each other; for instance, not every ’6’ is followed by a ‘7’ (but see Supplementary Material 6.14 for how this assumption can be relaxed). For each image, multiple inference-learning cycles (Eq 1-4) were conducted before switching to the next image in the sequence. A training epoch consisted of an iteration through all sequences from the dataset. Supplementary Material 6.2 contains the pseudocode for the nested training loops.

## 3. Results

We trained the network on sequences of moving objects as specified in the Methods section, and focused on the evolving high-level representations, resulting neural dynamics, and generative input-reconstructing capacities of the network, all in comparison to neurobiology.

### 3.1 Transformation-invariant stimulus representations

We found that neurons in network area 3 became tuned to samples in a position-invariant manner. To quantify invariance, we analyzed the neural representations in the highest area of trained networks (Fig 2) under changes of inputs. More specifically, inference was run on still images from the training datasets until convergence was reached (see Supplementary Material 6.3 for a description of convergence). Then, pairwise comparison of inferred area 3 representations measured in cosine distance quantified representational dissimilarity between representations of the same sample, e.g. a digit, (within-sequence) or different samples (across-sequence). All pairwise values were plotted in Representational Dissimilarity Matrices (RDMs, (57)) in Fig 4.

**Fig 4.**
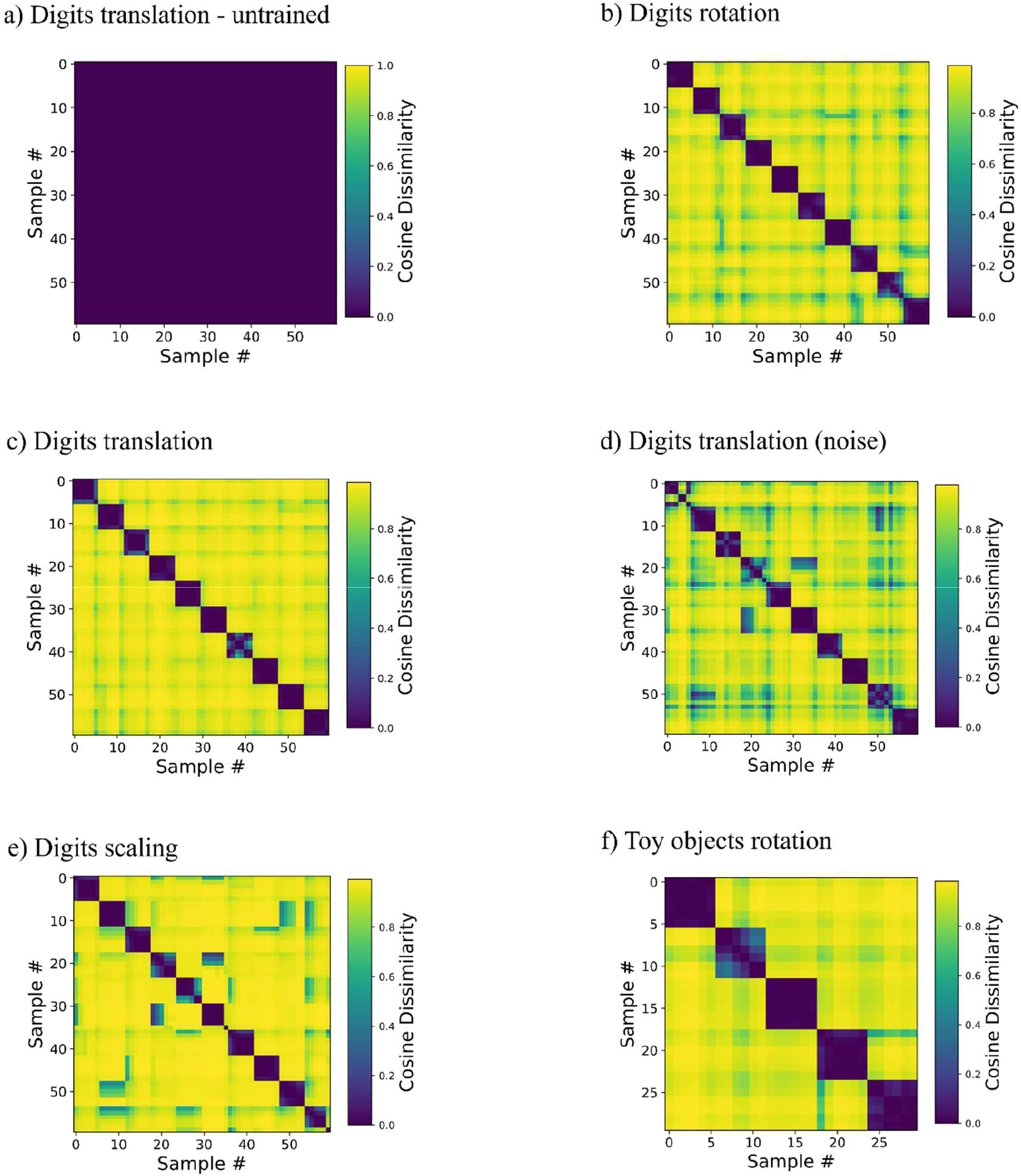
Representations invariant to viewing conditions are learned without data labels. The matrices depict cosine dissimilarity between representations in area 3. Each of the rows and columns in these plots corresponds to one input image (i.e. a digit sample in a specific spatial configuration), thus each matrix is symmetrical. Along each dimension, samples are ordered sequence-wise, i.e., rows and columns 0-5, 6-11 etc. are the same object in six different transformation states. Low values shown in purple correspond to similar activity patterns, i.e. a similar set of neurons represents the stimuli given by the combination of row and column, high values shown in yellow correspond to orthogonal activity vectors. a) Baseline, an untrained network tested on the translationally moving digits dataset, for untrained versions of the other RDMs see Fig S8. b)-e) Networks trained and tested on one of the three datasets of ten rotating, translating (with and without noise) and scaling digits show a clear block-diagonal structure with low values for comparisons within sequences. f) Network trained and tested on five rotation sequences of toy objects.

Indicating invariance, RDMs of trained networks showed high similarity within sequences; for instance, digit ”1” was represented by highly similar activity patterns in area 3, irrespective of position. Representations of samples from different sequences, such as digit ”1” and digit ”2” at the same position were distinct, as indicated by a high dissimilarity in matrix elements off the block diagonal. The same held true for the rotating and scaling digits (Fig4b and Fig 4e) as well as for the five rotating toy objects (Fig 4f). Supplementary Material 6.9 contains a proof of principle demonstration of learning multiple transformations in the same network.

Noise (shown for the translational motion in Fig 4d versus the noiseless motion in Fig 4c) slightly degraded clarity of the RDM but preserved the overall structure well. Additionally, the structure of the RDM proved to be quite tolerant to smaller weight initialization (Supplementary Material 6.7).

Invariance of representations was a consequence of learning from temporally continuously transforming inputs as evidenced by the RDMs of the untrained network that showed very little structure (Fig 4a, note the different color scale, cosine distance below 0.001). Networks trained on the static frames of the sequences, in which activity was reset after each frame also lacked a block-diagonal structure (Fig S2), illustrating the role of continuous motion in the training paradigm, which is to provide the necessary temporal structure in which subsequent inputs can be assumed to be caused by the same objects. Interestingly, we did not find an influence of sequence order on decoding accuracy (Fig S11), suggesting that only temporal (shown by the comparison to the static training paradigm), but not spatial continuity of the input transformations was necessary for successful representation learning. The Hebbian learning rule thus groups together consecutive inputs in a manner reminiscent of contrastive, self-supervised methods (35,36,59) that explicitly penalize dissimilarity in the loss function. Here, the higher-level representation from the previous timestep provides a target for the consecutive inputs reminiscent of implementations of supervised learning with local learning rules (53,60,61).

Area 3-representations were informative about the identity of the sample moving in sequence as decodability improved with training (Fig 5a-b). In addition to its behavioral relevance, decodability of representations quantifies the learned within-sequence invariance. A biologically plausible way to make high-level object representations available to downstream processes (such as action selection, Fig 1) is a layer of weighted synaptic connections, i.e., a linear decoder, to infer object identity. We simulated this through a linear mapping of the converged area 3-activity vectors that were obtained as above to ten object identity-encoding neurons (digits ”0”, ”1”, …, ”9”). After fitting the decoding model to 2/3 of the representations, evaluation was conducted on the remaining 1/3 in a stratified k-fold manner (with k = 3). Compared to the information content in the input signal, as measured by the accuracy of a linear decoder, as well as k-means clustering, area 3 representations achieved better decoding performance after around five training epochs (Fig 5a). The model also outperformed linear Slow Feature Analysis (SFA) (40) of the raw inputs (for details see Supplementary Material 6.12). This was confirmed across almost all used datasets (Tab 1) and even increased as the transformation step size was increased, resulting in smaller overlap between consecutive images (“fast” conditions in Tab 1, shown in the first and third row of Fig 3). Across the hierarchy, higher network areas developed more invariant representations than lower areas (Fig S9).

**Tab 1.** Decoding accuracy (in percent) across datasets and models. The best performing decoder per dataset is marked in bold. The left column is the predictive coding network trained in the continuous manner put forward in this paper.

| Dataset | PC-continuous,<br>area 3 | PC-static,<br>area 3 | k-means<br>input | Linear<br>decoding<br>input | SFA input |
| --- | --- | --- | --- | --- | --- |
| Toy objects rot. | **95.83** ±<br>**5.46** | 18.33 ±<br>1.67 | 53.33 | 83.33 | 73.33 |
| Digits rotating | **94.17** ±<br>**2.50** | 13.33 ±<br>2.64 | 48.33 | 85.00 | 70.00 |
| Digits rotating,<br>noise | **90.00** ±<br>**5.14** | 15.83 ±<br>1.86 | N/A | N/A | N/A |
| Digits scaling | 91.67 ±<br>2.64 | 75.83 ±<br>1.44 | 68.33 | **100.00** | **70.00** |
| Digits scaling,<br>noise | **86.67** ±<br>**4.25** | 71.25 ±<br>12.33 | N/A | N/A | N/A |
| Digits<br>translation | **83.75** ±<br>**6.50** | 40.42 ±<br>4.31 | 60.00 | 75.00 | 70.00 |
| Digits<br>translation,<br>noise | **93.33** ±<br>**4.08** | 47.92 ±<br>6.60 | N/A | N/A | N/A |
| Digits fast<br>rotation | **94.17** ±<br>**2.50** | 13.33 ±<br>2.64 | 31.67 | 40.00 | 70.00 |
| Digits fast<br>translation | **94.58** ±<br>**3.80** | 12.92 ±<br>0.72 | 38.33 | 25.00 | 70.00 |
| Digits fast<br>translation,<br>noise | **87.50** ±<br>**4.49** | 14.17 ±<br>3.00 | N/A | N/A | N/A |

**Fig 5.**
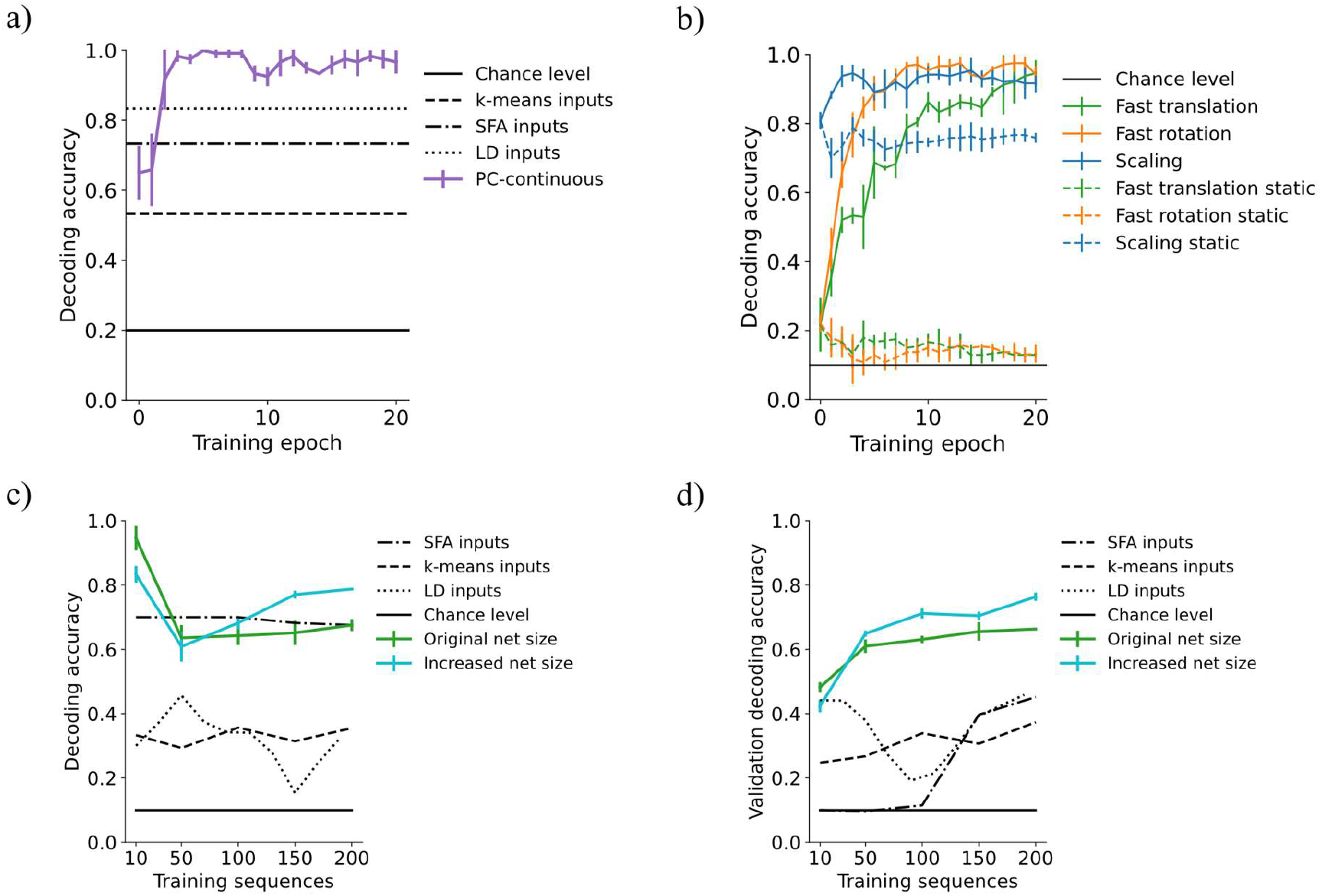
Area 3-representations encode object identity. Decoding accuracy of a linear decoder operating on area 3-representations of our predictive coding network trained in the continuous paradigm (PC-continuous) plotted across training epochs (iterations through the whole dataset). a) Accuracy quickly rises above performance of k-means clustering, SFA and linear decoding directly on the input data (LD inputs) for the rotating toy objects dataset. The error bars for all figures are computed across four random seeds for the weight initializations. b) Influence of continuous training: Decoding accuracy in networks trained on continuous sequences (continuous lines) is increased compared to networks trained on isolated (static) frames of the sequences. c) When increasing the size of the dataset from 10 to 200 sequences, the network of original size maintains a decoding accuracy far above chance level. Here, accuracy is significantly improved when the number of neurons in [area 1, area 2, area 3] is increased from [2000, 500, 30] (green curve) to [4000, 2000, 90] neurons (blue curve). d) Decoder accuracy on a previously unseen validation set of 200 randomly selected and transformed digits.

Decodability of network representations was maintained when the dataset size was significantly increased. We tested this by training networks on up to 20 random digits per digit class (totaling 200 sequences of the fast translations). As shown in Fig 5c, the network maintained above 60% linear decoding accuracy of digit class while an enlarged version of the network shown in cyan further improved this. On the other hand, increasing dataset size negatively affected the invariance structure of the RDMs (Supplementary Material 6.8). putatively due to the limitations discussed in section 4.3.

Lastly, generalization performance as measured by decoding accuracy on previously unseen digits was above 60% when more than 100 training sequences were used (Fig 5d).In the enlarged network, decoding accuracy rose above 75% (the blue line in Fig 5d), confirming the network’s capacity to generalize.

The continuous training paradigm improved decoding performance in comparison to networks trained on static inputs. There, decoding performance dropped from the initial value and was consistently more than twenty percentage points worse than in the continuously trained network (Fig 5b and Tab 1). This can partially be explained by the learning of more sample-specific and thus less invariant representations in the static training paradigm, where activity was not carried over from one image to the next (Fig S2).

### 3.2 Temporal stability of representations

Without explicitly integrated constraints, the network developed a hierarchy of timescales in which representations in higher network areas decorrelated more slowly over inference time than in lower areas. We quantified this by measuring the autocorrelation R during presentation of rotating digits. It is defined as

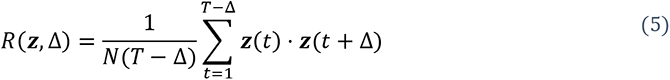

where Δ is the time lag measured in inference steps between the points to be compared, T is the duration of each sequence (consisting of 6000 inference steps), N the number of neurons in the subpopulation and z(t) the activity vector in the subpopulation (averaged across 10 inference steps). High values indicate similar, non-zero activities and thus high temporal stability. The resulting autocorrelation curves for time lags between 0 and the length of an individual sequence are shown in Fig 6a-b, averaged across the ten rotation sequences. From these curves, decay constants were inferred by measuring the time until decay to 1/e. If that value was not reached until the sequence end, we extrapolated by using a linear continuation through the values at Δ=0 and Δ=6000 time steps. The resulting decay times are shown in Fig 6c. A significant difference was found between representation neurons in area 3 and area 1 (mean difference: 7151 time steps, p=1.58e-8, determined by ANOVA), as well as a smaller, but still significant difference between representation neurons in area 2 and area 1 (4680 time steps, p=6.08e-6). In error-coding neurons, the hierarchy was less pronounced, but area 0 and area 2 nonetheless showed a significant difference (p=0.03). Comparison to a statically trained network with the same architecture which failed to develop a temporal hierarchy in representations (Supplementary Material 6.4) showed that the temporal hierarchy was not built into the model architecture, but instead is an emergent property of the model under the continuous training paradigm. This is underlined by the fact that the same inference rate was used in all network areas. The hierarchy in representational dynamics is in agreement with experimental findings in rat visual cortex (62). There, the authors computed neuronal timescales for the decay of autocorrelation in a similar manner and found more stable activity patterns in higher areas of rat visual cortex (Fig 6d).

**Fig 6.**
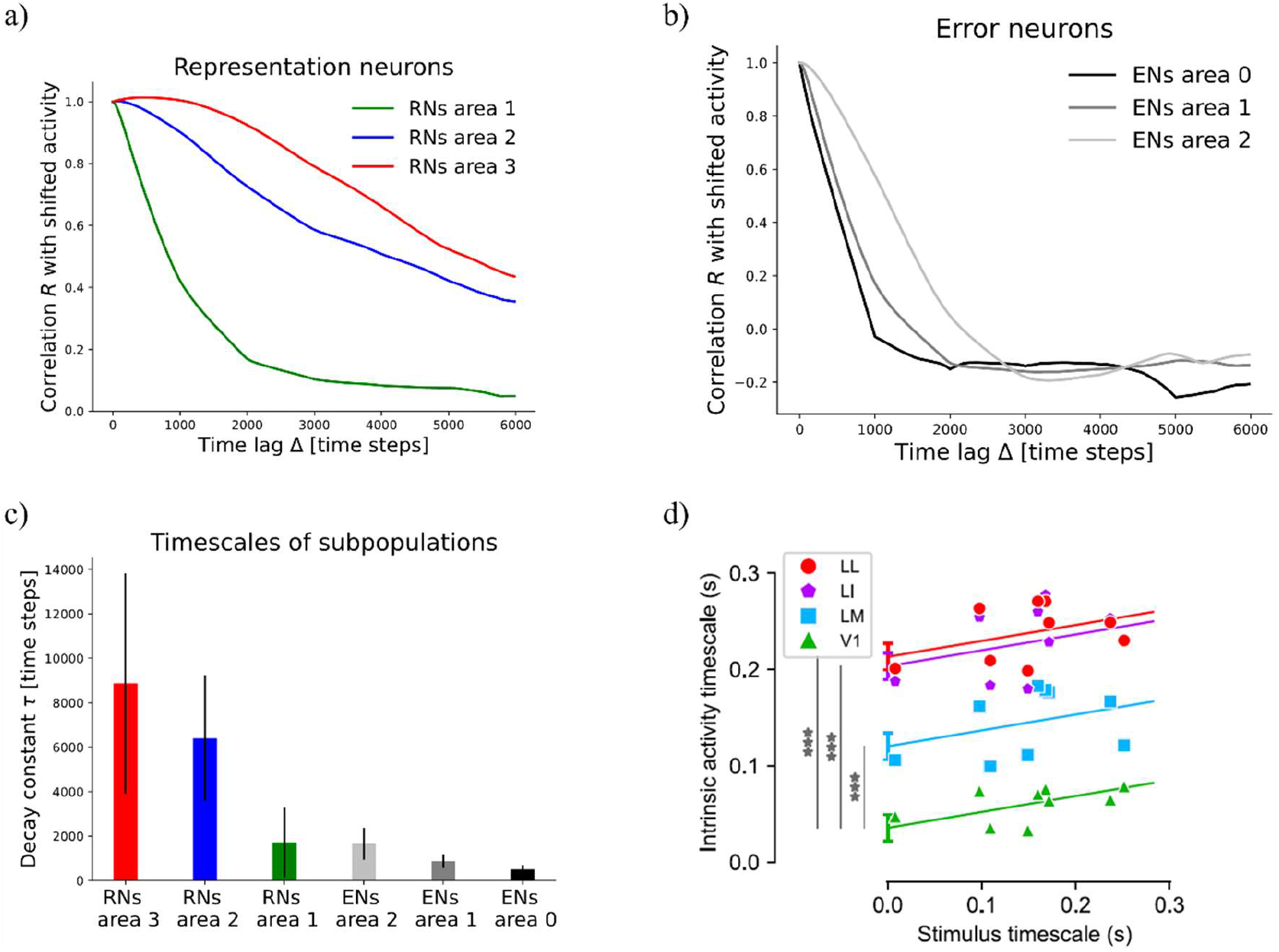
The network develops a hierarchy of timescales comparable to experimental data from rodent visual cortex. a) Temporal decay of activity autocorrelation for the representation neuron subpopulations (RNs). b) Decay of autocorrelation for the error-coding subpopulations (ENs). c) Inferred decay constants per subpopulation, p-values are given in the main text. d) Comparison to experimental evidence from rat visual cortex. Hierarchical ordering of intrinsic timescales rendered as the decay constants of activity autocorrelation, from V1 across lateromedial (LM), laterointermediate (LI) to laterolateral (LL) visual areas, adapted from (62). *** p=5e-7, 1e- 13, 2e-14, respectively for LM, LI and LL.

The decay speed of autocorrelation also allowed us to differentiate between quickly decorrelating error neurons and more persistent representation neurons in higher network areas. Error-coding neurons in area 2 showed a shorter activity timescale than representation neurons within the same area. The difference equaled 4740 time steps (p=3.9e-11), compared to only 211 time steps (p=2.87e-11) in the statically trained network (Fig S3). In this context, Piasini et al. (62) discussed the following scenario: When perceiving a continuously moving object, its identity is predictable over time. Thus, one could expect a diminishing firing rate in neurons representing this object, in contrast to their evidence on larger timescales in higher visual areas. Our results reconcile the framework of predictive coding with these empirical observations by differentiating between quickly decorrelating error-signals and persistent representations. Remarkably, this prediction about the consequences of predictive coding circuitry for the activity autocorrelation timescales of error- and representation neurons has, to our knowledge, not been proposed before. Here it is important to mention the extensive literature on the analysis of different frequency bands in cortical feedforward and feedback signal propagation (summarized in (63) from a predictive coding perspective). These sources did, however, not speak about temporal stability and the two concepts are not easily connected. It is, for instance, conceivable to have low-frequency signals that quickly decorrelate or high- frequency signals that are maintained over time.

### 3.3 Generative capacity

The network learned a generative model of the visual inputs as shown by successful input- reconstruction through the network’s top-down pathway (Fig 7). Since areas further up in the ventral processing stream of the cerebral cortex are thought to encode object identity, it is interesting to ask, in how far they are to be able to encode fully detailed scene information, or whether they contain only reduced information (such as object identity). To examine the functioning of this reverse pathway under the continuous transformation training paradigm, we investigated the representational content in each area by reconstructing sensory inputs in a top-down manner. After training, a static input image was presented until network activity converged (Supplementary Material 6.3). Then, the input was blanked out and the inferred activity pattern (representation) from a selected area was propagated back down to the input neurons via the top-down weights. Area by area, activity ***y***_l_ of representation neurons was installed by the descending predictions ***y****_l_* (see Supplementary Material 6.5 for details).

**Fig 7.**
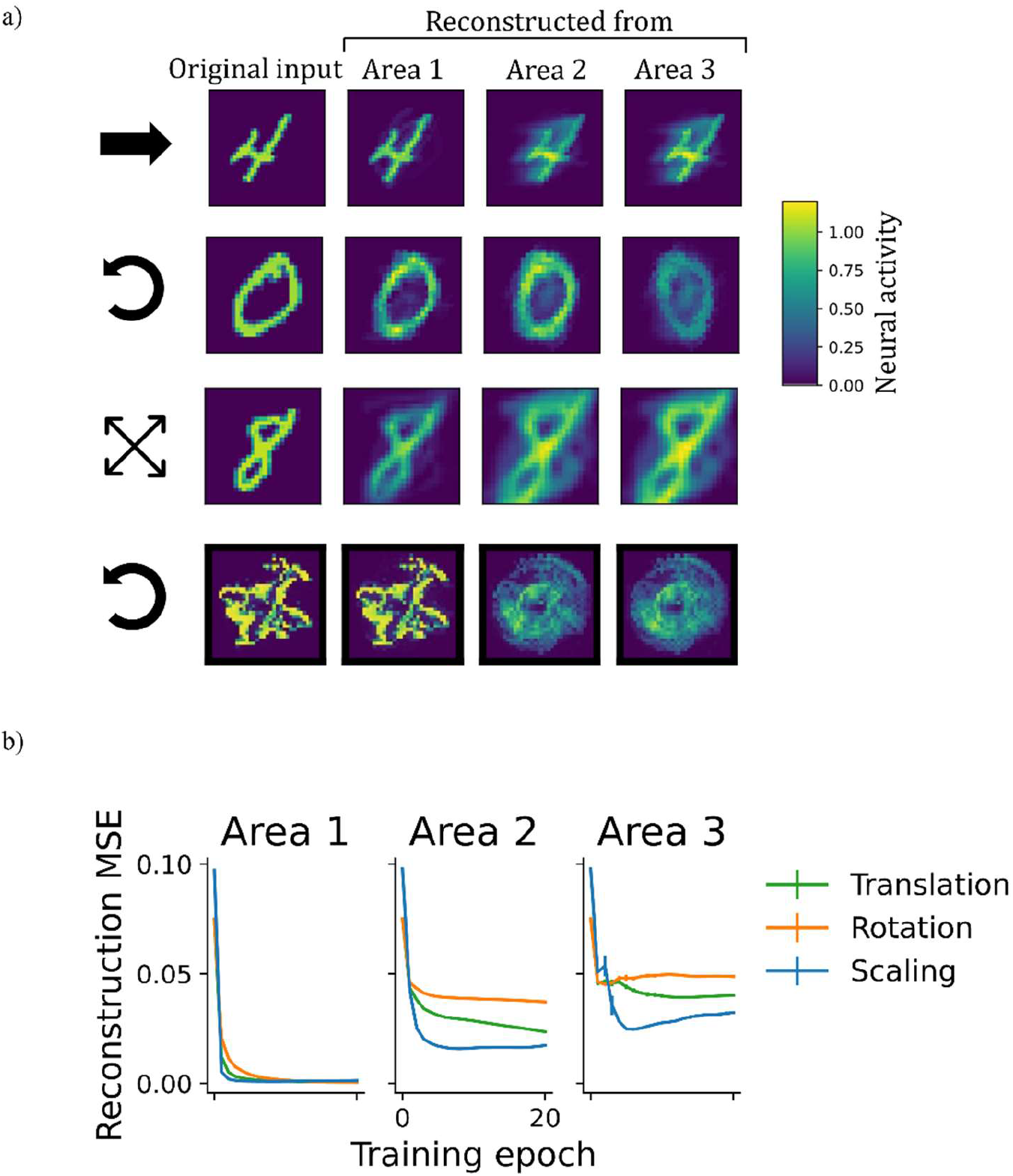
Learning of a generative model. a) Illustration of top-down reconstructions. The first column depicts original input images from different datasets. Columns two to four show the activity pattern in the input area generated by propagating latent representations from different network areas to the input layer in a top-down manner. In early network areas, representations inferred from sensory inputs carried enough information to reconstruct the input image once it was removed. Reconstructions from higher areas were less accurate. b) Mean squared reconstruction errors (MSE), comparing the original input to the reconstructions on a pixel-level. The vanishingly small vertical bars indicate the standard deviation across four random seeds.

As shown in Fig 7a, the accuracy of reconstruction strongly depended on the area it was initiated from. While predictions from latent representations in area 1 gave rise to reconstructions that resembled the original inputs and achieved low reconstruction errors (Fig 7b), higher areas were less accurate. From there, reconstructions were either blurry or showed the stimulus in a different position, rotational angle, or scale than presented prior to construction (e.g. the ”0” from area 3 in the second row of Fig 7). This logically follows from the invariance achieved in these higher areas, from where a single generalized representation cannot suffice to regenerate many specific images. Despite this limitation in obtaining precise reconstructions, which resulted from training on extended sequences instead of individual frames, area 1-representation neurons in all networks contained enough information to regenerate the inputs, thus confirming that the model had learned a generative model of the dataset.

### 3.4 Reconstructing objects from occluded scenes

The generative capacity of the network’s top-down pathway was further confirmed by its ability to reconstruct whole objects from partially occluded sequences as shown in Fig 8. A behaviorally relevant use of a generative pathway is the ability to fill in for missing information, such as when guessing what the whole scene may look like and planning an action towards occluded parts of an object. To investigate filling-in in the model, we presented occluded test sequences to the network trained on laterally moving digits (the same as before). After inference on each frame of the test dataset, the predictions sent down to the lowest network area were normalized and plotted retinotopically in Fig 8. Details on the reconstruction process can be found in Supplementary Material 6.6. Indeed, predictions sent towards the lowest area carried information about the occluded parts (Fig 8a).

**Fig 8.**
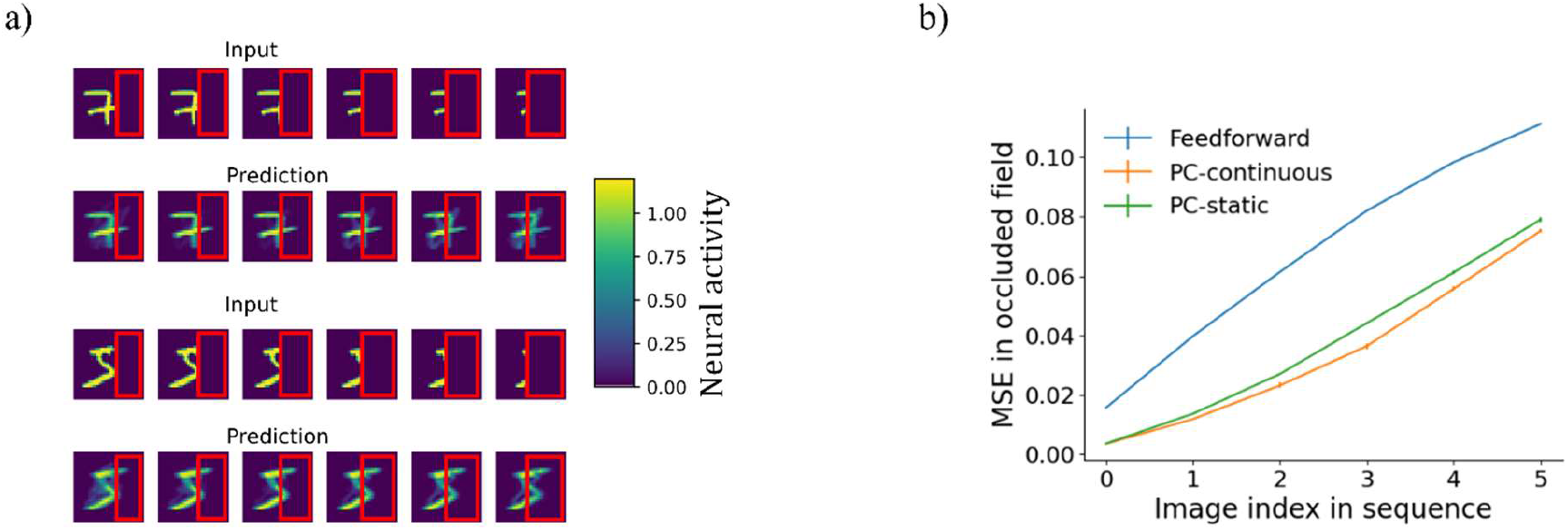
Reconstruction of partially occluded sequences. a) First and third row: the input sequences shown to the PC-continuous network with the occluder outlined in red. Rows two and four: images arising from top-down predictions sent to the area 0 carried information about occluded areas of the input. b) Comparison of a continuously trained predictive coding network to a purely feedforward network (no reconstruction) and a predictive coding network trained on static images. Shown is the mean squared reconstruction error in the occluded part, averaged across all ten sequences, rising as the occluded field becomes larger (plotted over the first to last image of the occlusion sequence). The vanishingly small error bars indicate the standard deviation across four network initializations.

As the input deteriorated, predictions also visibly degraded, resulting in a rising MSE (Fig 8b). The continuously trained network consistently achieved slightly, but significantly better reconstructions than its counterpart trained on static images (for a more detailed analysis see Supplementary Material 6.6). An independent t-test resulted in p<6e-4 for all sequence frames except for the first, unoccluded frame where the difference was non-significant. That the difference was small can be explained by two opposing mechanisms: on the one hand memorization of specific frames putatively aids reconstruction in the static network (Fig S2). On the other hand, availability of invariant object identity from the temporal context, which can be expected to improve reconstruction in the continuously trained network. Overall, the availability of top-down information in occluded fields of network area 0 is comparable to the presence of concealed scene information observed in early visual areas of humans (13) that cannot be explained by purely feedforward models of perception. Unlike auto-associative models of sequential pattern-completion (64), our network forms hierarchical representations comparable to (36).

## 4. Discussion

### 4.1 Summary of results

We have shown how networks that minimize local prediction errors learn object representations invariant to the precise viewing conditions in higher network areas (Fig 4), while acquiring a generative model in which especially lower areas are able to reconstruct specific inputs (Fig 7, Fig 8). The learned high-level representations distinguish between different objects, as linear decoding accuracy of object identity was high (Fig 5). Comparison to considerably worse decoding performance in networks trained on static images underlined the importance of temporally continuous transformation for the learning process (Fig 5a). Focusing on the implications for neural dynamics, learning from temporally continuous transformations such as continuous motion led to a hierarchy of timescales in representation neurons that showed more slowly changing activity in higher areas, where they notably differed from the more quickly varying error neurons (Fig 6).

### 4.2 A generative model to learn invariant representations

Without the need for explicit data labels, the model developed meaningful, decodable representations purely by Hebbian learning. Linking slowly varying predictions in higher areas to more quickly changing inputs in lower areas lead to emergence of temporally stable representations without the need for an explicit constraint for slowness as used for example in (40). At the same time, the model acquired generative capacity that enables reconstruction of partially occluded stimuli, in line with retinotopic and content-carrying feedback connections to V1 ((13, 38), see also (3) for a review of predictive feedback mechanisms). Other neuron-level models of invariance-learning (28,29,31,35) neither account for such feedback nor experimentally observed explicit encoding of mismatch between prediction and observation (10, 37) and used considerably more complex learning rules requiring a larger set of assumptions (35). Conversely, auto-associative Hopfield-type models that learn dynamic pattern completion from local learning rules (64, 65) do not learn hierarchical invariant representations like the proposed model does. By solving the task of invariance learning in agreement with the generativity of sensory cortical systems, the claim for predictive coding circuits as fundamental building blocks of the brain’s perceptual pathways is strengthened. We argue that the model generalizes predictive coding to moving stimuli in a biologically more plausible way than other approaches (16,46,50) that rely on non-local error backpropagation (44) or backpropagation through time (50). While the neural circuitry of our model is similar to previous implementations of predictive coding with local learning rules (5,18,53), the consequences of training such networks on dynamic inputs (invariance, neuronal dynamics) had not been investigated so far. Biologically, dispensation of a term depending on the partial derivative with respect to neuronal activity minimizes the set of necessary assumptions compared to other implementations that require such a term in inference (18, 53) or learning (18). Unlike (18), the present implementation also does not require weight regularization that depends on information not readily available at the synapses.

### 4.3 Limitations in performance

Although sufficient for learning of invariant representations on small datasets, the fully connected architecture we used can be expected to limit the degree of representation invariance (visible e.g. in the structure of the RDMs) on larger datasets. Fully connected areas may also restrict performance on out-of-sample testing. Here, combination of receptive field-like local filters with a pooling mechanism (66) may be helpful to become tolerant to the varying configurations of individual features comprising the objects from the same class. Using a weakly supervised paradigm could improve decoding accuracy even further. It has been shown that under constraints which would be out of the scope of this paper to discuss, predictive coding can do exact backpropagation when clamping the highest layer activities in a supervised manner (53, 67).

Input reconstructions from higher network areas degraded as representations became more invariant. This is a direct consequence of Eq 6: Each element from the set of area 3- representations casts a unique prediction to the area below. Consequently, multiple different (not invariant) area 3 patterns would be necessary to fully reconstruct a sequence of inputs. Thus, either the invariance in area 3 or the faithfulness of the reconstruction suffers. Nevertheless, the network as a whole appeared to strike a good balance in the trade-off of memorizing information to reconstruct individual samples in lower areas (hence the better reconstruction accuracy from area 1 in Fig 7) and abstracting over the sequence, where area 3 represents object identity invariantly (Fig 5), fitting theoretical descriptions of multilevel perception (ch. 9 in (19)). The more detailed and sample-specific information may provide useful input to the action-oriented dorsal processing stream (68), whereas the hierarchy of the ventral visual cortex extracts object identity and relevant concepts (69).

### 4.4 Hypotheses on the neural circuitry of predictive coding

The model captures neural response properties in early and high-level areas of the visual cortical hierarchy. Retinotopic (38) and information carrying (13) feedback to early visual areas (cf. Fig 7; Fig 8) as well as invariant (24, 25) and object-specific representations (cf. Fig 4) in the temporal lobe (21–23) are captured by the simulation results. While there is ample evidence for a hierarchy of timescales in the visual processing streams of humans (70), primates (71) and rodents (62), with larger temporal stability in higher areas, the compatibility with deep predictive coding is debated (62). Our simulation results of increasingly large timescales further up in the network hierarchy may help to reconcile predictive coding with the experimental evidence. Coincidentally, this was also found to be true in a recently developed predictive coding model, albeit with only two layers and without explicit error representations (43). Compared to emergence of temporal hierarchies purely as a result of dynamics in spiking neurons (72) or large-scale models (73, 74), our model provides a complementary account, postulating development of the temporal hierarchy as a consequence of a functional computation: learning invariance by local error minimization.

What novel insights can be extracted about the brain’s putative use of predictive algorithms? Theories of predictive coding range from limiting it to a few functions (such as subtraction of corollary discharges to compensate for self-motion (10)) and input reconstruction (5) to claiming extended versions of it as the most important organizational principle of the brain (4), namely the free energy principle. PC models provide a critical step to make theories of perception and imagery quantitative and falsifiable as well as to guide experimental research (3). Based on the simulation results, error neurons in higher visual areas operate on a much shorter activity timescale than their representational counterparts. This comparison of distinct subpopulations may provide an additional angle to measuring neural correlates of prediction errors (for a review see (75)), as representation neuron responses have been barely considered in experimental work so far. In combination with work on encoding of errors in superficial, and representations in deep cortical layers (3,63,76,77), area- and layer-wise recordings of characteristic timescales could lead to a better understanding of cortical microcircuits underlying predictive coding. Layer-wise investigations also show distinct patterns of feedforward and feedback connectivity (78) and information processing (79). Only with knowledge about these microcircuits, models of finer granularity can be constructed.

### 4.5 Conclusion

Predictive coding is a theory with great explanatory power, but with unclear scope. Here, we go beyond the original scope of pure input-reconstruction and find that predictive coding networks can additionally solve an important computational problem of vision. Our results are in line with experimental data from multiple species, strengthening predictive coding as a fundamental theory of mammalian perception.

## Acknowledgements

The authors would like to thank Shirin Dora and Kwangjun Lee for constructive discussions.

This project has received funding from the European Union’s Horizon 2020 Framework Programme for Research and Innovation under the Specific Grant Agreement No. 945539 (Human Brain Project SGA3; to C.P. and S.B.).

We acknowledge the use of Fenix Infrastructure resources, which are partially funded from the European Union’s Horizon 2020 research and innovation program through the ICEI project under the grant agreement No. 800858.

## 6. Supplementary Materials

### 6.1 Network architecture and hyperparameters

The networks consist of four fully connected areas of which the topmost (area 3) has no error units and the lowest (area 0) is driven by the input only, in a pixel-wise manner. Thus, the number of neurons in area 0 corresponds to the resolution of the inputs. Tab S1 shows the precise number of neurons per area and simulation. Tab S2 lists the used hyperparameters.

**Table S1.** Network architecture used for the different datasets.

| Dataset | Architecture |
| --- | --- |
| Translation MNIST | 1156, 2000, 500, 30 |
| Rotating MNIST, Scaling MNIST | 784, 2000, 500, 30 |
| Toy objects | 1156, 1000, 250, 15 |

**Table S2.**
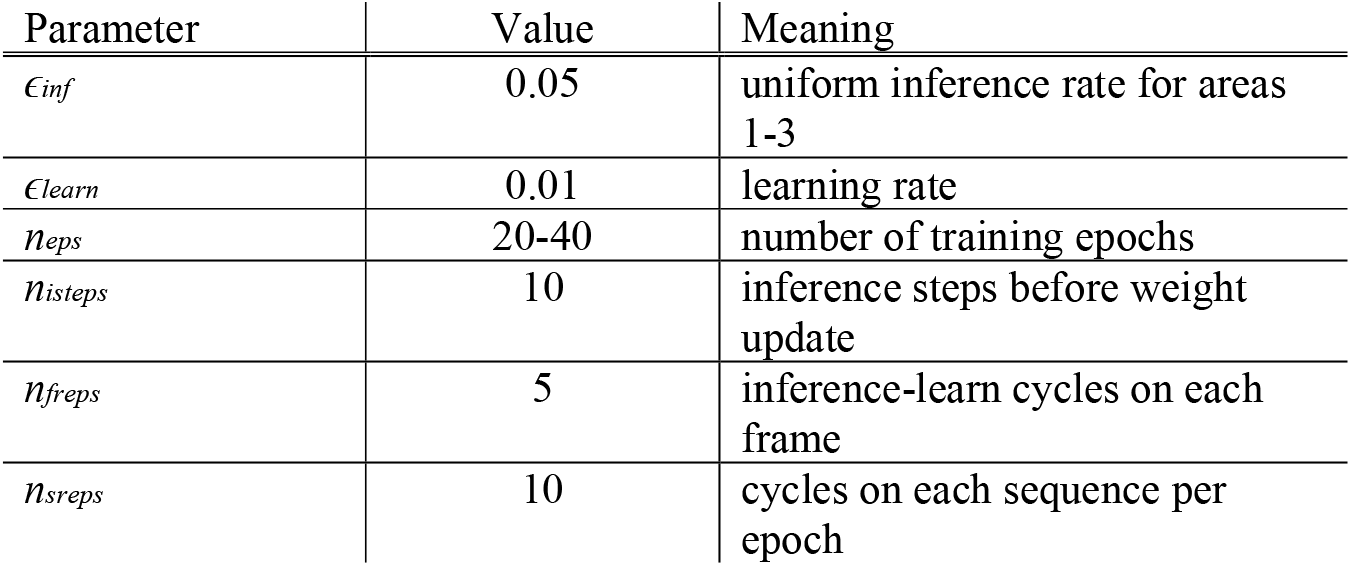
Network hyperparameters.

### 6.2 Training

Network training was conducted in nested loops as described in section 2.3. Alg S1 depicts the pseudocode for both training paradigms. The main difference between the continuous and static training paradigms is the additional reset of activity in the beginning of the fifth ‘for’ loop. To compensate for the shorter time between activity resets in the static training paradigm, the number of cycles per frame *n_freps_* is increased by a factor of 6.

**Alg S1.**
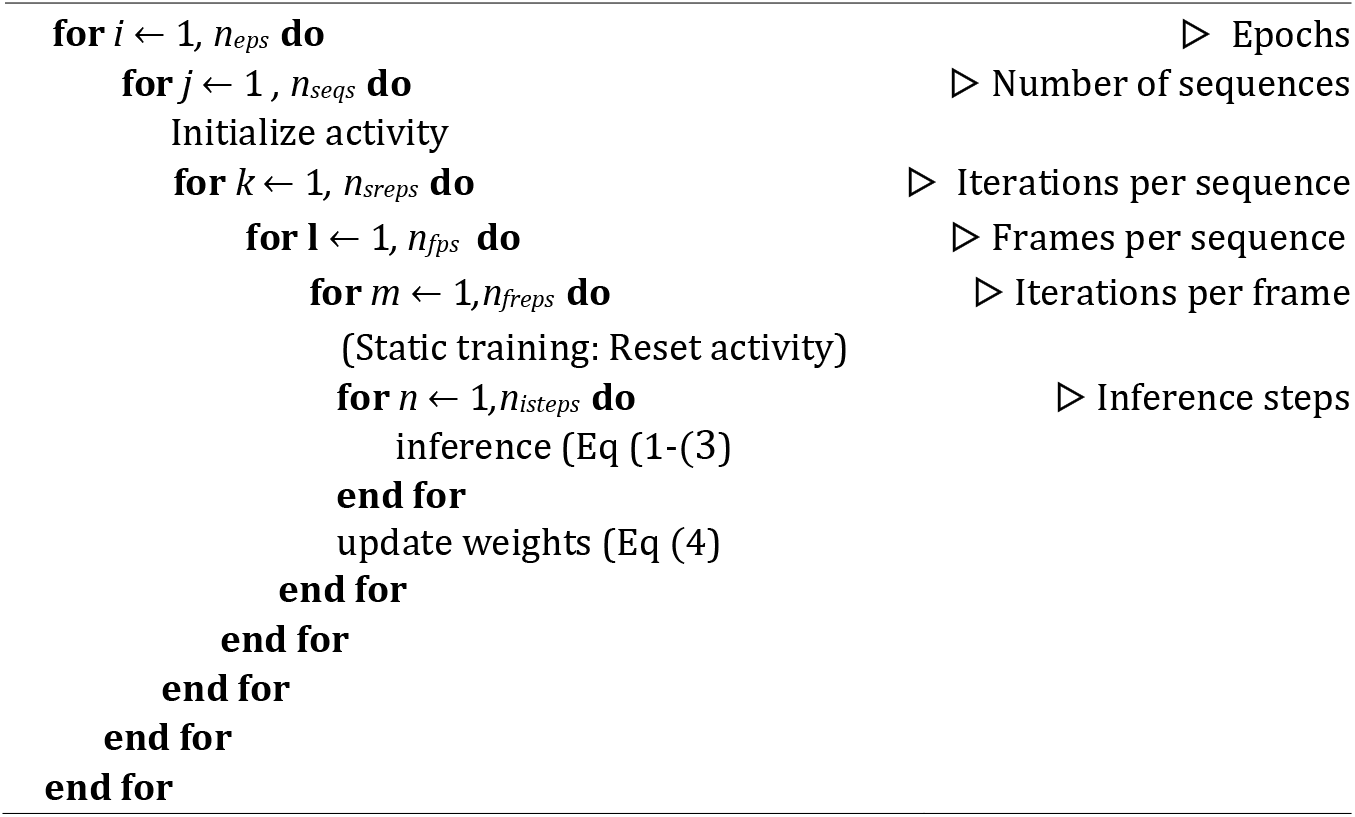
Pseudocode for network training. Nested loops during network training as described in the Methods section. In the static training, activities are reset each time a new input is presented.

### 6.3 Inference of high-level representations

Convergence of high-level neural activity took considerably longer (2000 inference steps were used) than the duration that was necessary until the total prediction error in the network converged (one to two orders of magnitude less). I.e., after the prediction errors were minimized, further inference improved invariance, but not stimulus reconstruction. Across the network areas, this is plotted for the rotating digits in Fig S1.

**Fig S1.**
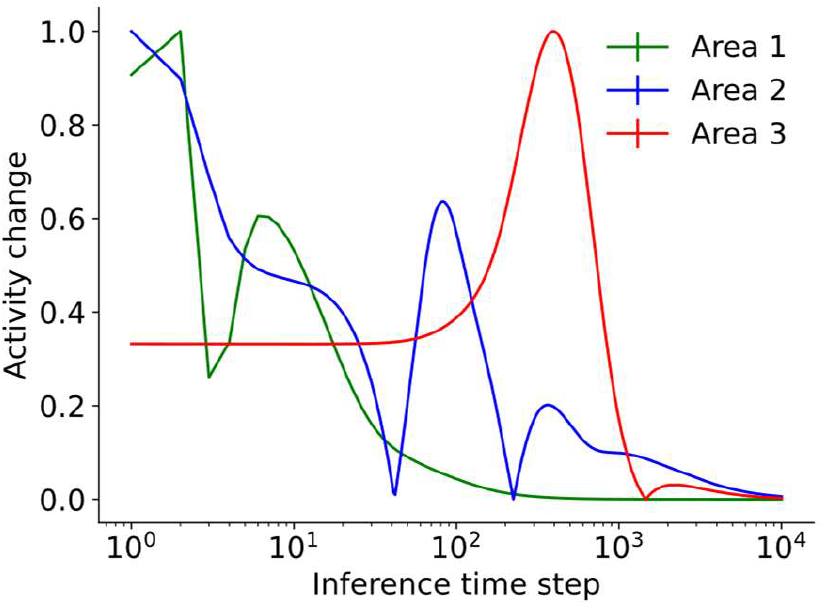
Convergence of activity over inference steps for a fixed inputs from the rotating digits dataset across representation neurons in the different network areas.

### 6.4 Influence of temporal continuity

In how far is the learning of invariant representations attributable to the training paradigm? Fig S2 shows the RDM of a network trained in a static manner: the more position-dependent representations stand in contrast to the RDMs of the continuously trained networks (Fig 4b-f) with more uniform representations within sequences.

To analyze whether the different neural timescales are a result of the training paradigm or a result of the network architecture, we compared the decay speed of activity autocorrelation to the statically trained network (Fig S3).

**Fig S2.**
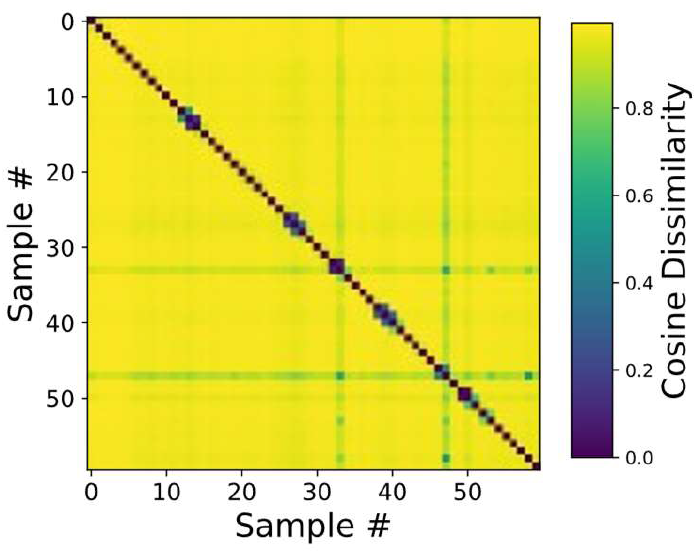
Influence of static training on high-level representational invariance. Network trained in a static manner with resets of activity after each frame show less invariance in area-3 representations compared to the continuously trained networks (Fig 4b-f). The diagonal structure of the RDM shown here reveals that representations are specific to individual images and do not generalize across positions.

**Fig S3.**
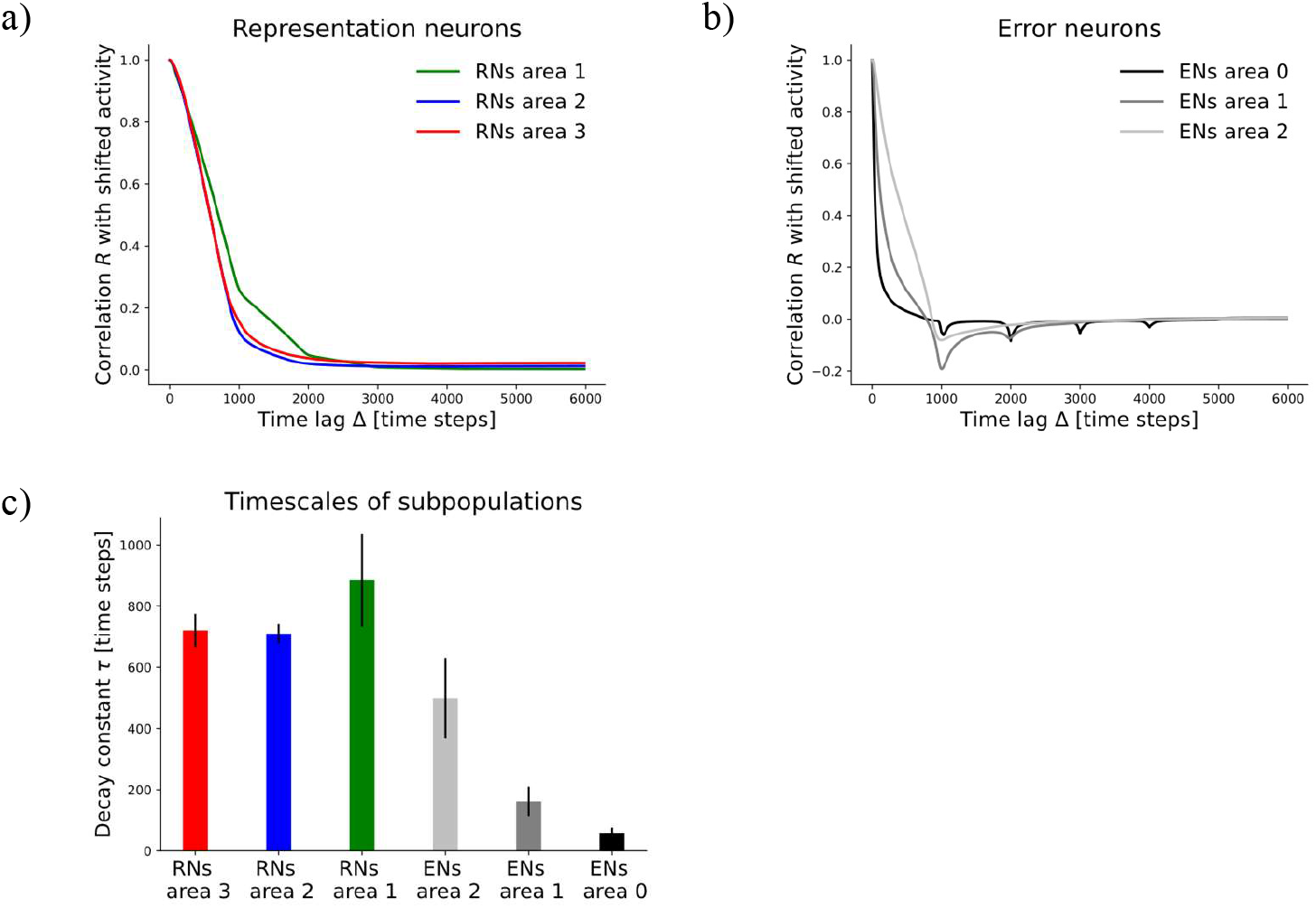
Decay of activity autocorrelation of neural subpopulations in a statically trained network in which activity is reset after each frame. a) Representation neurons, b) Error neurons, c) Inferred decay constants to quantify the timescale on which neural activity changed. This figure can be compared to Fig 6, but note the different scale in Fig 6c.Reconstruction of complete inputs

To test which input patterns can be reconstructed from different network areas, an input image needs to be presented for an extended period. Here, 2000 inference steps were used. The image was then removed, and reconstructions were triggered from the chosen area, i.e., predictions were consecutively projected to the area below according to Fig 2a:

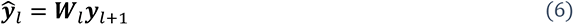

For symbols, the reader is referred to Eq 1.

### 6.5 Reconstruction of occluded sequences

For reconstruction of whole objects from partially occluded sequences, the first frame of a sequence was presented for an initial number of inference steps (see below) to give higher areas time to infer stimulus identity. Each consecutive frame of the sequence was then shown for a varying number of consecutive inference steps that is specified later. Lastly, reconstructions were normalized, dividing all predictions by the value of the largest pixel-wise prediction of the current reconstruction. To systematically investigate the difference between the static and continuous training paradigm, we varied both the initial and consecutive number of inference steps across a range of parameters and used the optimal combination we found for Fig 8. The continuously trained network consistently achieved better reconstructions than the statically trained network as shown by the consistently positive mean difference in reconstruction errors in Fig S4.

**Fig S4.**
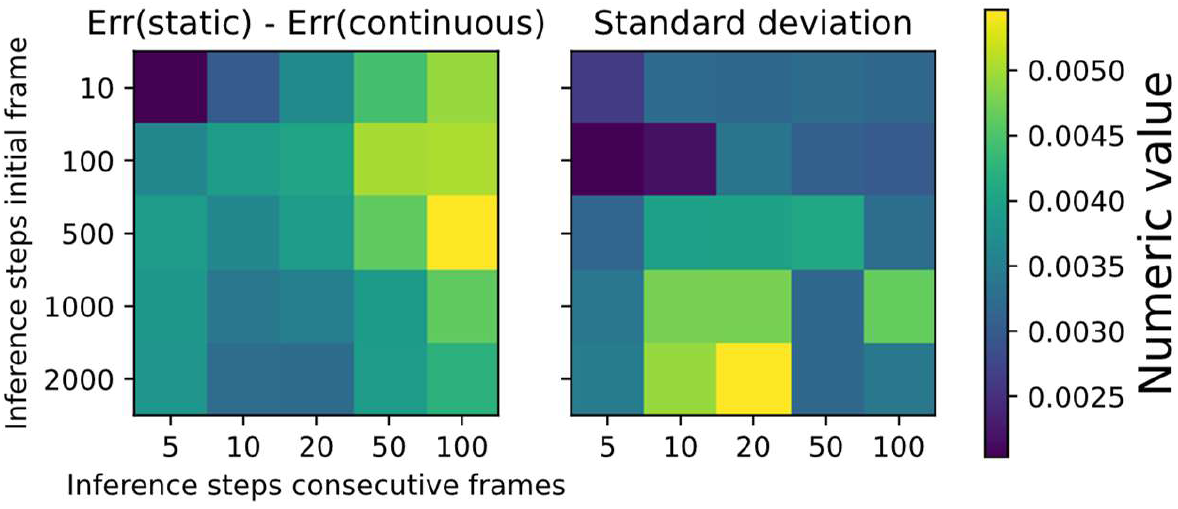
Reconstruction of occluded scenes: influence of hyperparameters. The continuously trained network achieved better reconstructions of occluded inputs than the statically trained one. The left plot shows the mean difference in reconstruction error between the two training paradigms across ten occlusion sequences, for various hyperparameters (see main text). Positive values indicate better performance of the continuously trained network. The right plot shows the standard deviations across the ten occlusion sequences.

### 6.6 Influence of weight initialization

Initializing weights to lower values than used by default (for default values see section 2.2) led to a decrease in performance, with less distinguishable representations in area 3 after 10 epochs (Fig S5). To test this, we decreased the standard deviation of the Gaussian weight initialization. As the distribution is centered at zero and clipped at zero to prevent negative weights, the value of the standard deviation corresponds to the average value of the initial weights (note that this is before dividing each weight by the number of neurons in the next higher area). Despite initially less distinguishable representations for smaller initial weights, prolonged training led to a non-collapsed state even for the smallest initialization that we tested (on the right of Fig S5).

**Fig S5.**
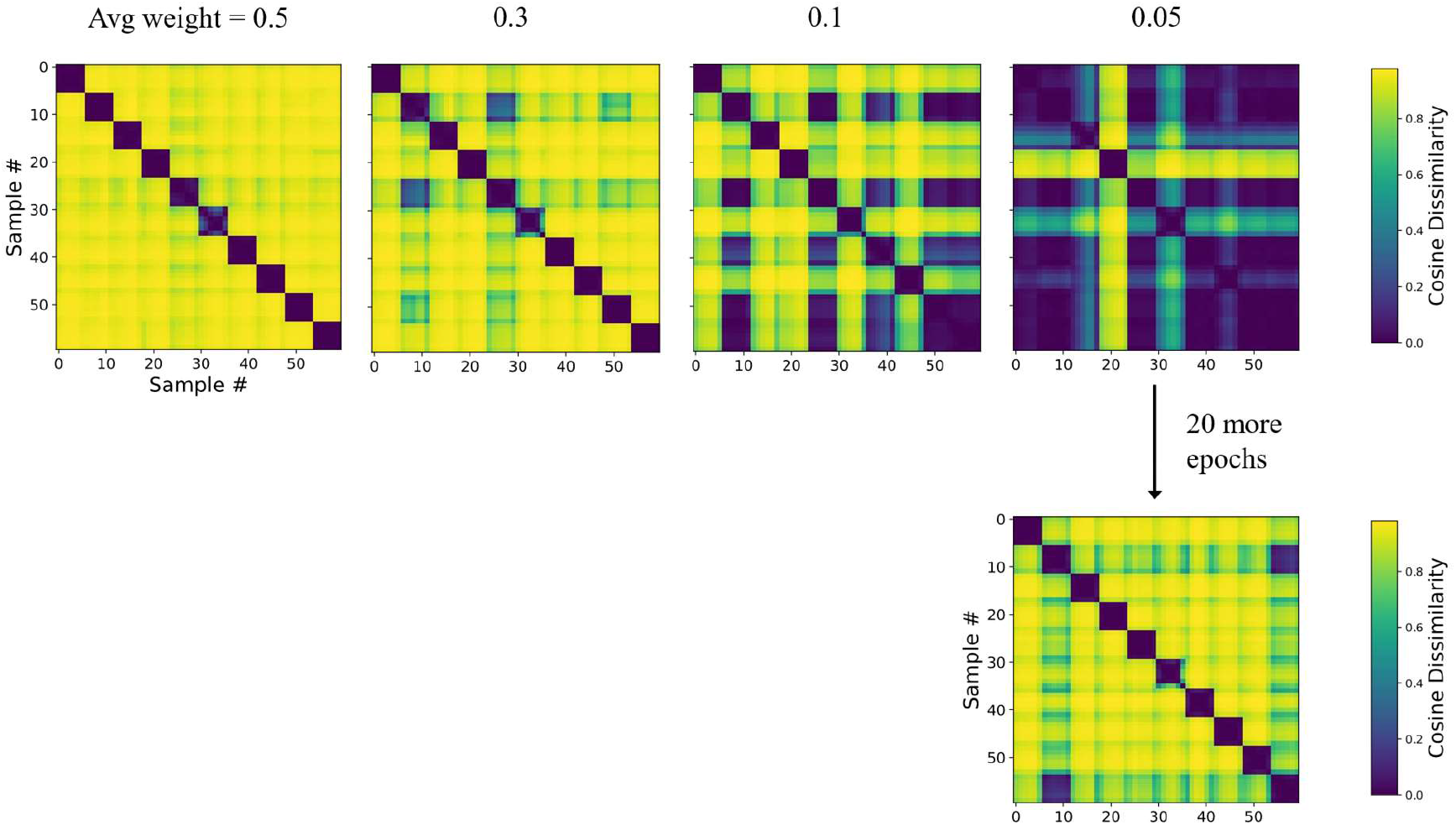
Influence of weight initialization. Initializing weights to smaller values gradually deteriorates the RDMs that are shown after 10 training epochs. The leftmost plot corresponds to the default settings used in the rest of the paper, shown for a digit translation dataset. The rightmost plot corresponds to a tenth of these weight values. After 20 more epochs on the network trained with an initial weight of 0.05, a considerable improvement was observed.

### 6.7 Representational dissimilarity matrices on larger datasets

To test the influence of increasing the dataset size on the RDMs, we ran the same analysis of constructing the RDMs (as in Fig 4) for the networks trained on the larger datasets. While the RDMs shown in Fig S6 lost the clear block-diagonal structure, decoding performance was still far above chance (Fig 5c). This effect, that was present for both the default and the increased network size, can be explained by a better approximation of the underlying data-generating distribution when dataset size was increased.

**Fig S6.**
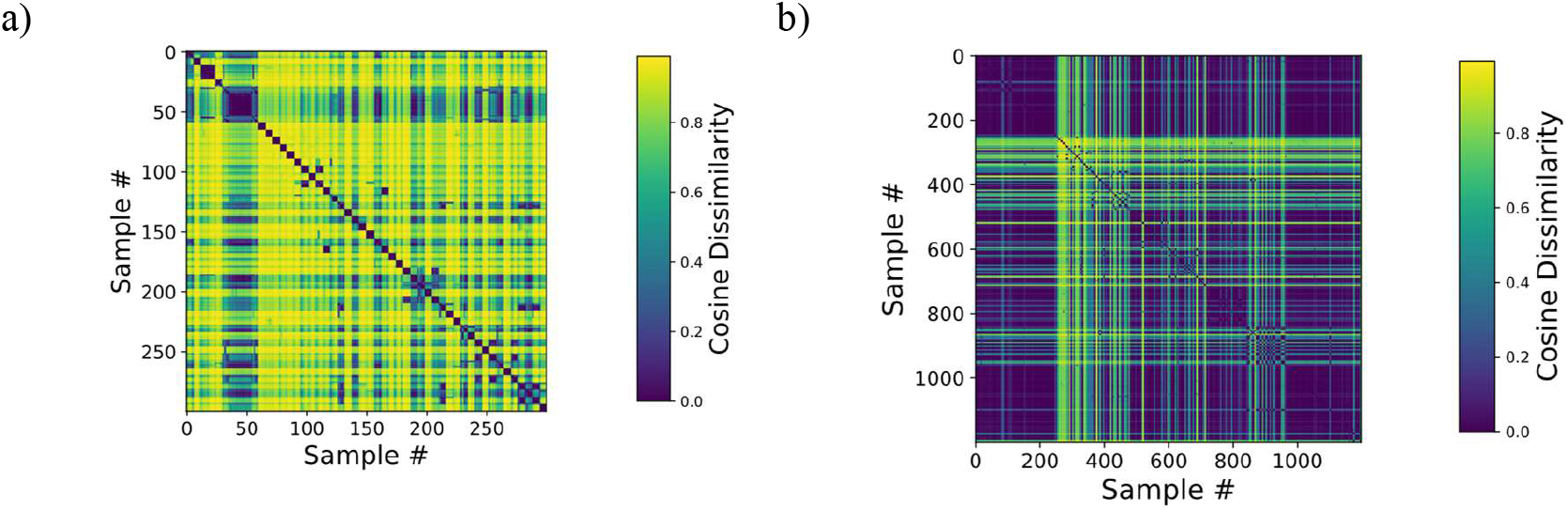
Representational dissimilarity matrices on larger datasets. When dataset size is increased such that multiple digits of each class are shown (in different sequences), the network forms instance-specific (e.g. a particular digit ‘1’ irrespective of position) instead of class-specific (e.g. all digit ‘1’s) representations. a) RDM for a network of increased size, with [4000, 2000, 90] neurons in [area 1, area 2, area 3] (chosen because it better visualizes the effect) instead of the default [2000, 500, 30]. The network was trained and evaluated on a dataset of 50 digit instances (sequences), five ‘1’s, five ‘2’s, etc., resulting in 300 image frames. The blocks along the diagonal are of size six, as each sequence contains six frames. b) When increasing the dataset size even further, the structure in the RDM becomes less obvious, but the digit classes are still relatively well decodable (Fig 5c). An explanation is given in the main text.

### 6.8 Learning multiple transformations in a single network

Learning different transformations of the same digit when these are not shown in sequence proved challenging. Nevertheless, Fig S7 provides a proof of principle result showing decoding of digit identity with ∼80% accuracy after seven training epochs, with a linear decoder again trained on 2/3 of the inferred representations. With prolonged training, the network tended to separate the invariant representations for the different transformations of each digit, suggesting the need for a unifying mechanism to merge these different invariant representations.

**Fig S7.**
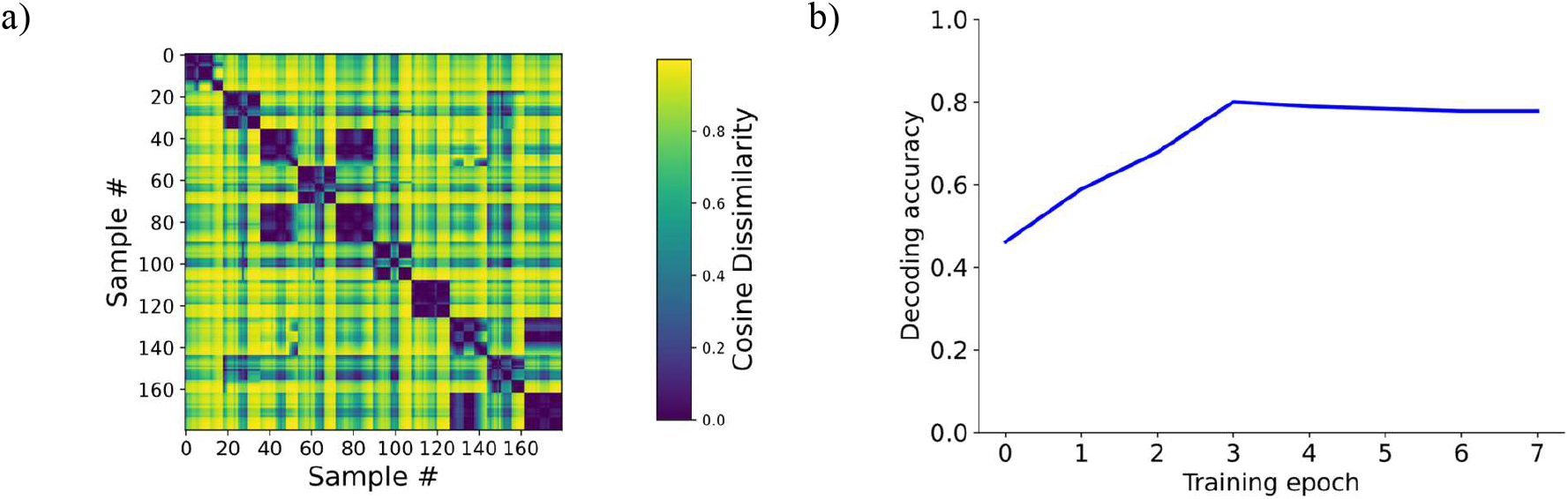
Learning multiple transformations in a single network with [2000,500,10] neurons in [area 1, area 2, area 3]. a) RDM of ten digits each undergoing three transformations (translation, rotation, scaling). b) Linear decoding accuracy of digit identity from area 3 of the same network.

### 6.9 Comparison of continuously trained to the untrained network

Complementing Fig 4, Fig S8 depicts the comparison of the continuously trained against the untrained network for the remaining datasets (listed in the figure caption).

**Fig S8.**
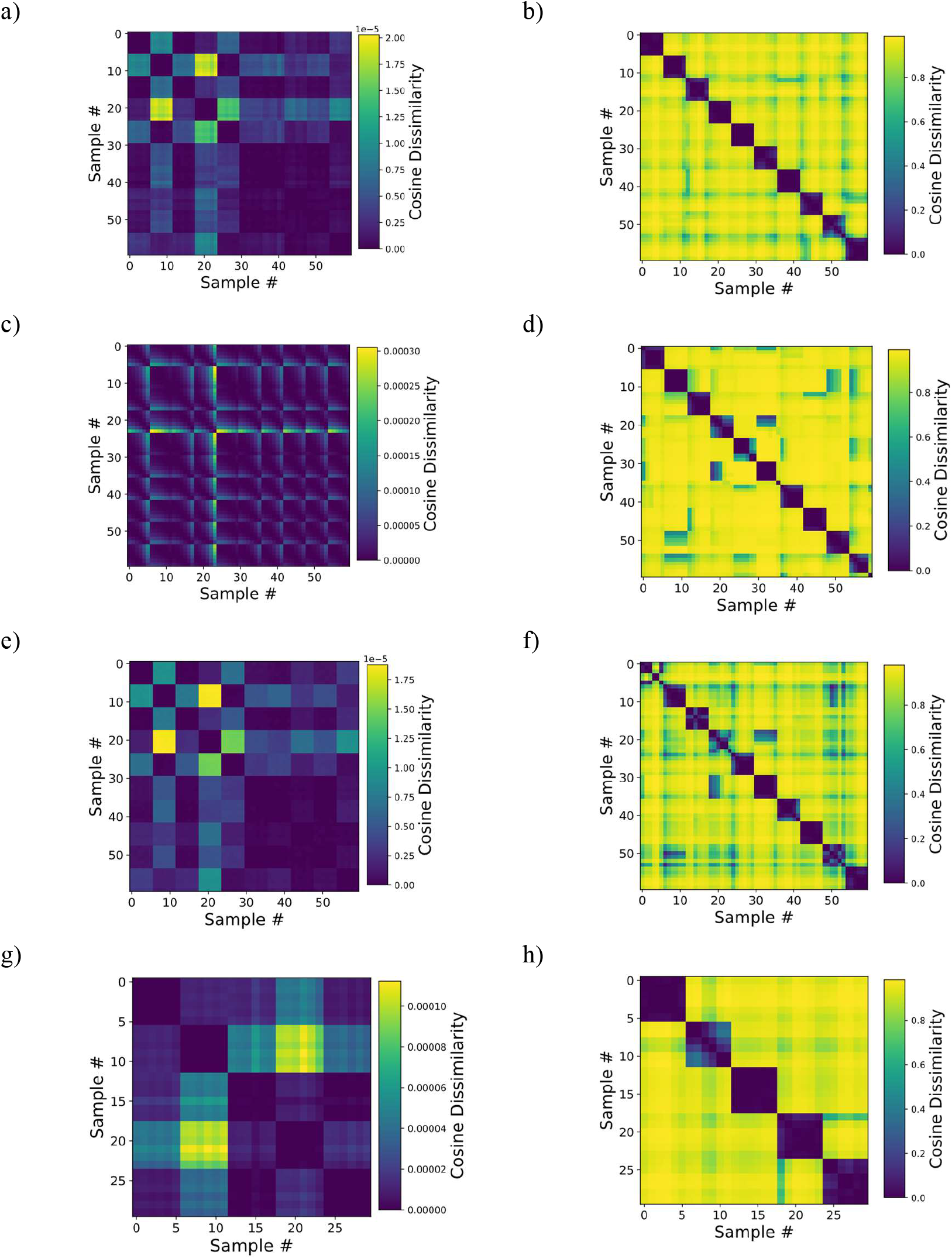
Comparison of continuously trained to the untrained network. Complement to Fig 4: comparison of untrained RDMs (left) to trained (right). a) and b) digit rotation, c) and d) digit scaling, e) and f) digit translation with noise, g) and h) rotating toy objects.

### 6.10 Comparison across areas

Higher network areas contained more invariant representations than lower areas. We quantified this by computing a sequence invariance value S, that is given as the ratio between the average cosine distance *d̄_in_* between representations of frames from the same sequence (after 2000

inference steps each) and the average distance to frames from all other sequences *d̄_across_*:

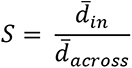

As shown in Fig S9, area 2 initially rises quickest, but is then surpassed by area 3. The result is related to the hierarchy of timescales, as more invariant representations (in areas with a higher value according to Fig S9) can be expected to be more stable across time.

**Fig S9.**
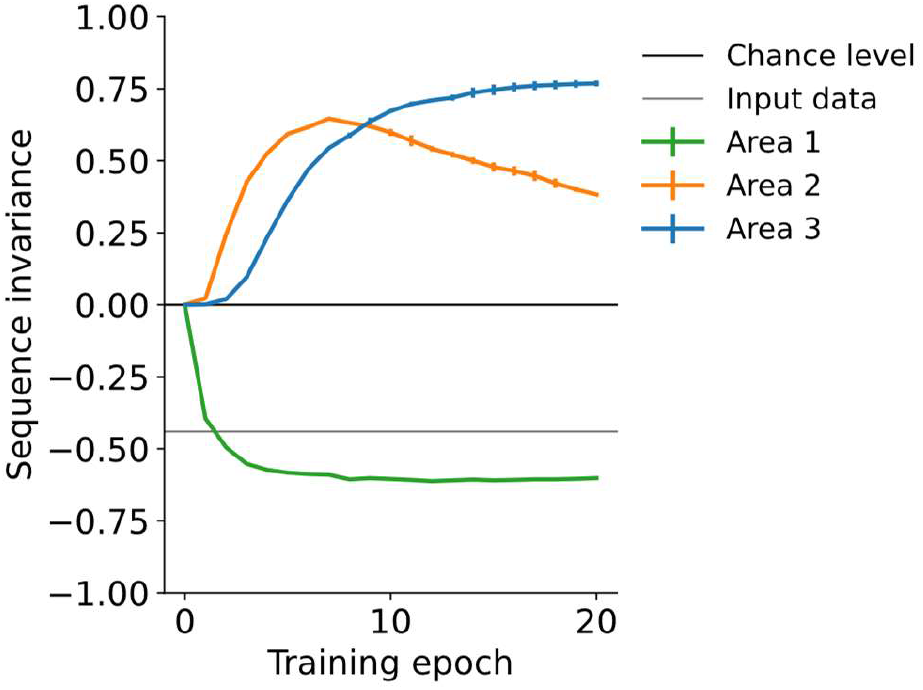
Comparison of representational invariance across network areas. The highest network area (area 3) eventually develops more invariant representations as compared to lower areas, which is mirrored in an eventually larger sequence invariance value. Error bars indicate standard deviation across two random seeds (computed on the fast translation digits).

### 6.11 Comparison to Slow Feature Analysis

To compare model performance to linear Slow Feature Analysis (SFA) (40), we used the sklearn-sfa package from https://pypi.org/project/sklearn-sfa/. From the concatenated digit/toy object input sequences, we let the algorithm extract a fixed number of features. As this number of features needs to be provided for the SFA algorithm, we tried different settings and optimized for decoding accuracy on each dataset. The subsequent linear decoding analysis consisted of training a linear decoder in a stratified k-fold manner on 2/3 of the extracted features and evaluated on the remaining 1/3. For the five moving digits, 30 features – matching the number of neurons in our top layer - proved optimal. For the five toy objects, reducing the number of features from 30 to 10 significantly improved accuracy from 23.33% to 73.33%, so we used this setting for the results shown in Fig5a and Tab 1.

### 6.12 Changing the number of neurons in different network areas

As it could be expected that projecting from the representation in one area to a larger number of neurons in the next higher area orthogonalizes the representations in the higher area, we systematically analyzed the influence of varying the size of all three network areas across almost two orders of magnitude. The results depicted in Fig S10 show the robustness of the learning paradigm and a minimum number of required neurons in area 3. As this analysis was performed on a dataset with ten samples only, we also refer to the simulations on larger datasets (Fig 5c-d) that showed slightly improved performance when network size was increased. Although orthogonalization may play a role, the memorization capacity of area 3 is more important here: the more sequences are shown, the more neurons are necessary to represent them separately.

**Fig S10.**
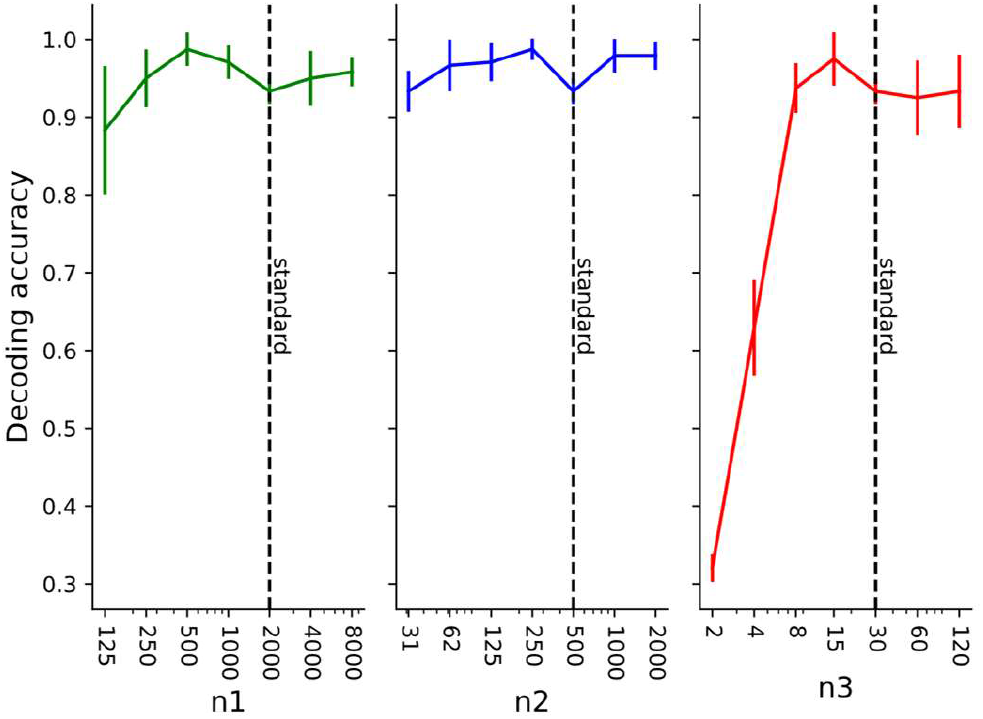
The effect of changing the number of neurons in the respective areas on decoding accuracy of object identity from area 3 representations. On the x-axis, n1 corresponds to the number of neurons in area 1, n2 and n3 accordingly in areas 2 and 3. The vertical “standard” line refers to the network architecture used in most simulations, except where noted differently.

### 6.13 Influence of sequence order

While influence of temporal continuity becomes clear when comparing to the static training paradigm (Fig 5b) it is not obvious whether only temporal proximity is necessary or whether the transformation also needs to be spatially continuous. To test this, we trained the network from Fig 5b on shuffled versions of the input sequences. We found that irrespective of sequence order, the only important factor was the temporal continuity of the transformation (Fig S11). In a less ethologically plausible paradigm that would differ significantly from the scope of this paper, this fact could be exploited by showing distinct instances from a class consequently without resets. In this case, the preselection of which images to show in sequence, however, would require labeled data and thus be supervised. Additionally, it should be noted that movement and spatial continuity are difficult concepts in fully connected networks without an a priori retinotopic layout.

**Fig S11.**
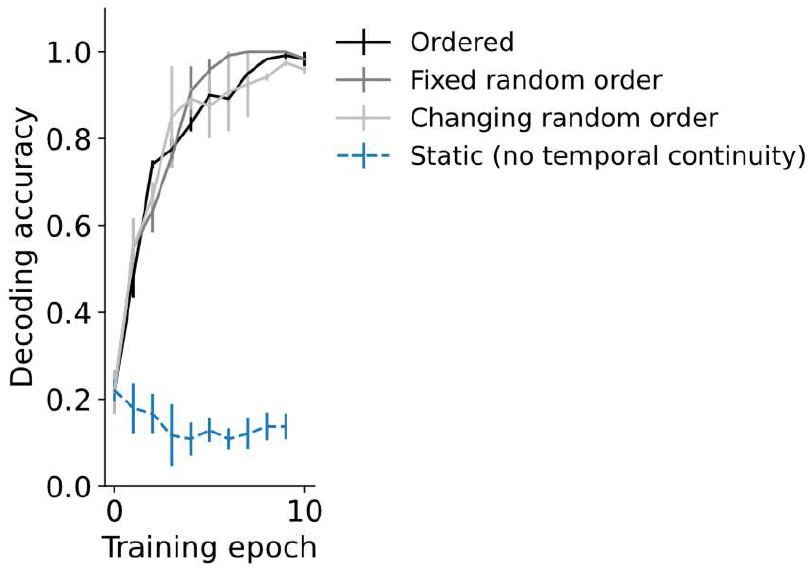
Temporal continuity but not sequence order influences decoding accuracy of object identity. Four conditions are compared: No temporal continuity (the static training paradigm) and three temporally continuous transformations. In addition to the spatially continuous transformation used in the main text (‘ordered’), sequences were presented in a fixed random order and in a changing random order that was shuffled newly each epoch.

### 6.14 Reducing the number of activity resets between sequences

To investigate how representation learning is influenced by not resetting the network activity at the start of each new sequence, we introduced a probabilistic reset. As shown in Fig S12a, resetting the activity approximately every three sequences slightly reduced the linear decodability. This decrease in performance follows from the less separated representational manifolds as shown in the RDMs (Fig 12b). Merging of representations in turn is caused by distinct digits being now presented in sequence with high probability (a ‘5’ has a 70% chance of being followed by a ‘6’ without activity reset). As mentioned in the main text, such a situation is rare in nature where objects are typically encountered in an interleaved manner, thus reducing the likelihood of merging their representations.

**Fig S12.**
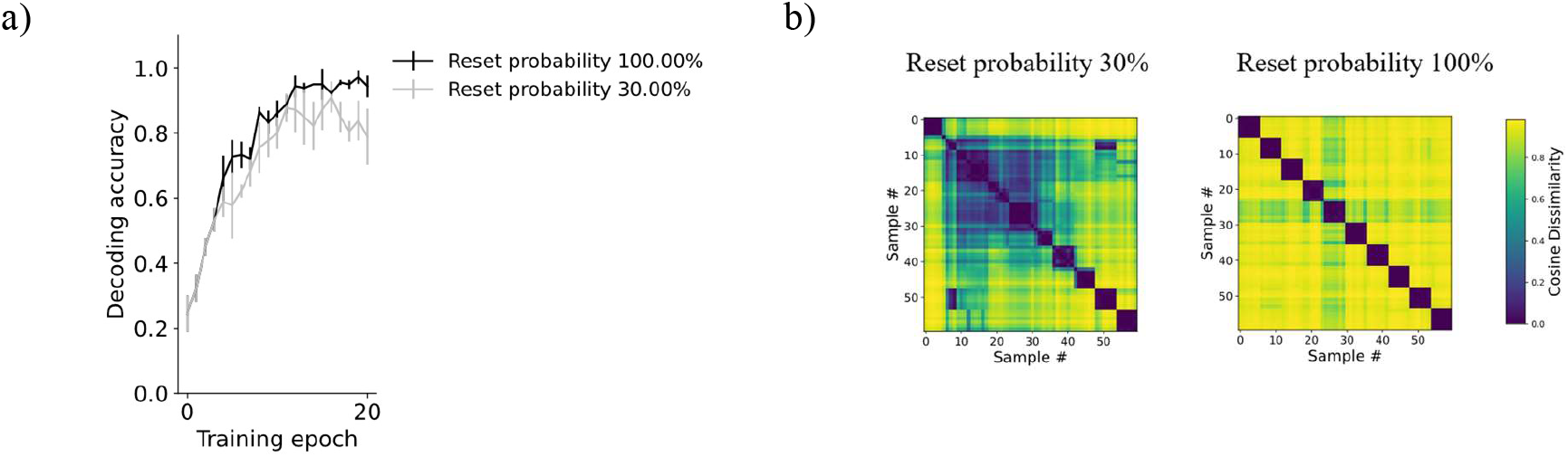
Reducing the number of activity resets maintains good representation learning. a) Linear decoding accuracy in the standard training paradigm and with reduced activity resets, shown for the fast digit rotation dataset. Small differences to the simulations in the main script (Fig 5b) result from sequential instead of batchwise training being deployed here. b) RDM for both conditions from a).

## Notes

### Competing Interest Statement

The authors have declared no competing interest.

### Summary of Updates

- Analyze the influence of sequence order - Add baseline to upscaling simulations - Analyze influence of reduced probability for activity reset

